# The EPS-I exopolysaccharide transforms *Ralstonia* wilt pathogen biofilms into viscoelastic fluids for rapid dissemination *in planta*

**DOI:** 10.1101/2025.05.22.655385

**Authors:** Matthew Cope-Arguello, Jiayu Li, Zachary Konkel, Nathalie Aoun, Tabitha Cowell, Nicholas Wagner, A. Li Han Chan, Lan Thanh Chu, Samantha Wang, Mariama D. Carter, Caitilyn Allen, Lindsay J. Caverly, Loan Bui, Kristen DeAngelis, Matthew J. Wargo, Tuan M. Tran, Jonathan M. Jacobs, Harishankar Manikantan, Tiffany M. Lowe-Power

**Author notes:** **Co-corresponding authors** Harishankar Manikantan and Tiffany M. Lowe-Power, **Email:** and.

## Abstract

*Ralstonia solanacearum* species complex (RSSC) pathogens cause destructive plant wilt diseases of a wide variety of crops, leading to significant agricultural losses worldwide. These bacteria rapidly spread through the water-transporting xylem where they grow prolifically and produce abundant biofilm that clogs xylem vessels. To understand RSSC biofilm behavior *in planta*, we examined their complex fluid mechanics. Rheological analyses revealed that unlike all previously analyzed microbial biofilms, RSSC biofilms are shear-thinning, viscoelastic fluids at physiologically relevant shear forces. To determine which factors confer these unique mechanics, we analyzed biofilms of bacterial mutants with altered biofilm components. Genetic analysis demonstrated that development of the viscous-dominant biofilms required production of EPS-I, an amphiphilic exopolysaccharide that is a major virulence factor for all RSSC pathogens. We show that EPS-I confers “biofilm mobility”, which allows wildtype RSSC colonies to passively expand when deformed. Despite its high metabolic cost, bioassays demonstrated that EPS-I production conferred a net fitness benefit where biofilm mobility allowed the pathogen to spread and access more nutrients in complex environments like xylem vessels. The RSSC are a monophyletic lineage of aggressive plant wilt pathogens, and our evolutionary hypothesis testing suggests the origin of the *eps* biosynthetic gene cluster coincides with the emergence of wilt pathogenesis in the RSSC ancestor. Furthermore, comparative physiological assays demonstrated that biofilm mobility is unique to the RSSC within the genus *Ralstonia*. In summary, EPS-I production is a key evolutionary innovation that enables RSSC dispersal and virulence by conferring unique biofilm mechanics.

**Significance Statement:** *Ralstonia solanacearum* species complex (RSSC) pathogens threaten global food security by fatally wilting plants. A soft matter physics lens demystified the cryptic role of a major virulence factor, the EPS-I exopolysaccharide. EPS-I transforms RSSC biofilms into viscoelastic fluids, a mechanical behavior not previously described for other microbial biofilms that are almost always viscoelastic solids. We demonstrate that the development of fluid biofilms was a key evolutionary innovation that enabled pathogenic success of these aggressive pathogens that rapidly wilt plants.

## Introduction

Approximately 80% of bacteria on earth live in complex, three-dimensional aggregates called biofilms (1). Biofilms are assemblages of one-or-more microbial species embedded in a self-produced matrix of extracellular DNA (exDNA), exopolysaccharides, lipids, and proteins. By developing biofilms, bacteria remodel their immediate physical, chemical, and biological environments. The biofilm environment confers resilience to environmental stresses, including antimicrobial molecules, desiccation, predation by phages, and attacks from microbial competitors.

Biofilms have adhesive, viscous, and elastic mechanical properties that enable many bacteria to form stable and persistent colonies in dynamic environments, including in plant and animal hosts (2). The mechanical behavior of diverse microbial biofilms has been extensively studied using rheometry (**Table S1**) (3–23). Although biofilm mechanics differ based on genetic and environmental variation, biofilms typically behave as viscoelastic solids, meaning they have a larger elastic component (termed *G`*, or elastic modulus) compared to the viscous component (termed *G*``, or viscous modulus). Conversely, viscoelastic fluids would have a greater viscous component than elastic. For viscoelastic materials such as a biofilm, acute, fast forces impose short-lived stresses, causing the biofilm to spring back like an elastic band (2). In contrast, slow, persistent forces relax polymeric molecular springs, causing the biofilm to flow like a fluid, enabling passive expansion of the biofilm. Thus, the mechanical properties of biofilms influence how bacteria colonize complex, dynamic environments, like those with flowing conditions.

Bacterial plant pathogens within the *Ralstonia* genus produce notoriously fluid biofilms *in planta* and on agar (24). Collectively, these causal agents of bacterial wilt disease are known as the *Ralstonia solanacearum* species complex (RSSC), a monophyletic lineage of three species that colonize the water-transporting xylem vessels and disrupt xylem sap flow across a broad range of hosts (25). Microscopy of RSSC-infected xylem tissues usually shows substantial biofilms of RSSC cells and extracellular matrix (26–29). These *in planta* biofilms allow diagnosticians to identify the tell-tale sign known as bacterial streaming (30), where visible plumes of pathogen biofilms “stream” out of xylem vessels when the cut stem of a wilted plant is submerged in water. RSSC pathogens develop biofilms by producing a matrix of extracellular DNA, proteins, and a unique amphiphilic, exopolysaccharide called “EPS-I” (29, 31, 32). These components contribute to the structure and function of RSSC biofilms. RSSC secrete two nucleases, NucA and NucB, which modulate the amount of extracellular DNA (29). Δ*nucAB* mutants form thicker biofilms and dome-shaped colonies, suggesting that DNA influences the mechanical behavior of RSSC biofilms. RSSC secrete multiple sugar-binding lectin proteins (32), and mutants lacking the LecF and LecX lectins develop expansive colonies with apparent zones of collapse in the center (32). However, the most important biofilm component is the major virulence factor EPS-I (29, 30, 33–35). RSSC pathogens secrete abundant EPS-I when growing at high cell densities *in vitro* and in plant hosts because the expression of the corresponding *eps* biosynthetic gene cluster is positively regulated by quorum sensing (36–40). Exactly how EPS-I contributes to bacterial wilt disease has perplexed researchers (24). A potential model is that EPS-I enables pathogen dissemination throughout the host (33, 41, 42).

Here, we hypothesize that RSSC pathogens have evolved to produce biofilms with unique mechanical properties, which contribute to pathogenic success by rapidly spreading the pathogens throughout the flowing xylem environment. To test this hypothesis, we employed rotational rheometry to quantify the bulk mechanical behavior of colony biofilms from diverse wildtype RSSC isolates, from RSSC mutants with altered biofilm composition, and from an unrelated plant pathogen. In striking contrast to all previously investigated bacterial biofilms that are viscoelastic *solids*, we show that EPS-I production transforms RSSC biofilms into viscoelastic *fluids*. This mechanical behavior enhances RSSC fitness by promoting colony expansion in both *in vitro* and *in planta* environments, highlighting the adaptive advantage conferred by fluidal biofilms. Our evolutionary genomic, physiological, and rheological analyses suggest that the emergence of the RSSC as rapid wilt pathogens coincided with the evolutionary invention of fluidal biofilms, offering new insight into the ancient emergence of these globally impactful pathogens.

## Results and Discussion

### RSSC wilt pathogens form biofilms that are shear-thinning, viscoelastic fluids

Peculiarly, cultures of RSSC isolates appear to be more liquid than solid (**Fig. 1A-B; Movie S1**), suggesting that their biofilms are not typical viscoelastic solids. We investigated the rheological properties of RSSC biofilms with selected representatives of each of the three RSSC species (*R. solanacearum* “*Rsol*” IBSBF1503, *R. pseudosolanacearum “Rpseu*” GMI1000, and *R. syzygii* “*Rsyz*” PSI07) (**Fig. 1C-D, Fig. 2, Fig. S1**, and **Fig. S2**). These isolates have subtle natural diversity in their colony morphology (**Fig. S3**). *Rpseu* GMI1000 and *Rsyz* PSI07 produce typical colonies that are wide, fluidal, and have irregular shape whereas *Rsol* IBSBF1503 produces smaller dome-like colonies characteristic of the “small, fluidal, round (SFR)” morphology (43, 44). These data revealed an inverse relationship between colony size and viscosity (**Fig. 1D** and **Fig. S3**). At a shear rate of 0.11 s^-1^, the RSSC biofilms had mean viscosities of 2.19 ± 0.639 Pa*s (*Rpseu* GMI1000), 6.47 ± 0.629 Pa*s (*Rsol* IBSBF1503), and 1.39 ± 0.160 Pa*s (*Rsyz* PSI07) (± standard deviation [SD]; **Fig. 1D**). To further understand the implications of these viscosities, we determined that this shear rate is relevant to the shear rates of xylem flow (**SI “Results and Discussion”**) (45).

**Fig. 1.**
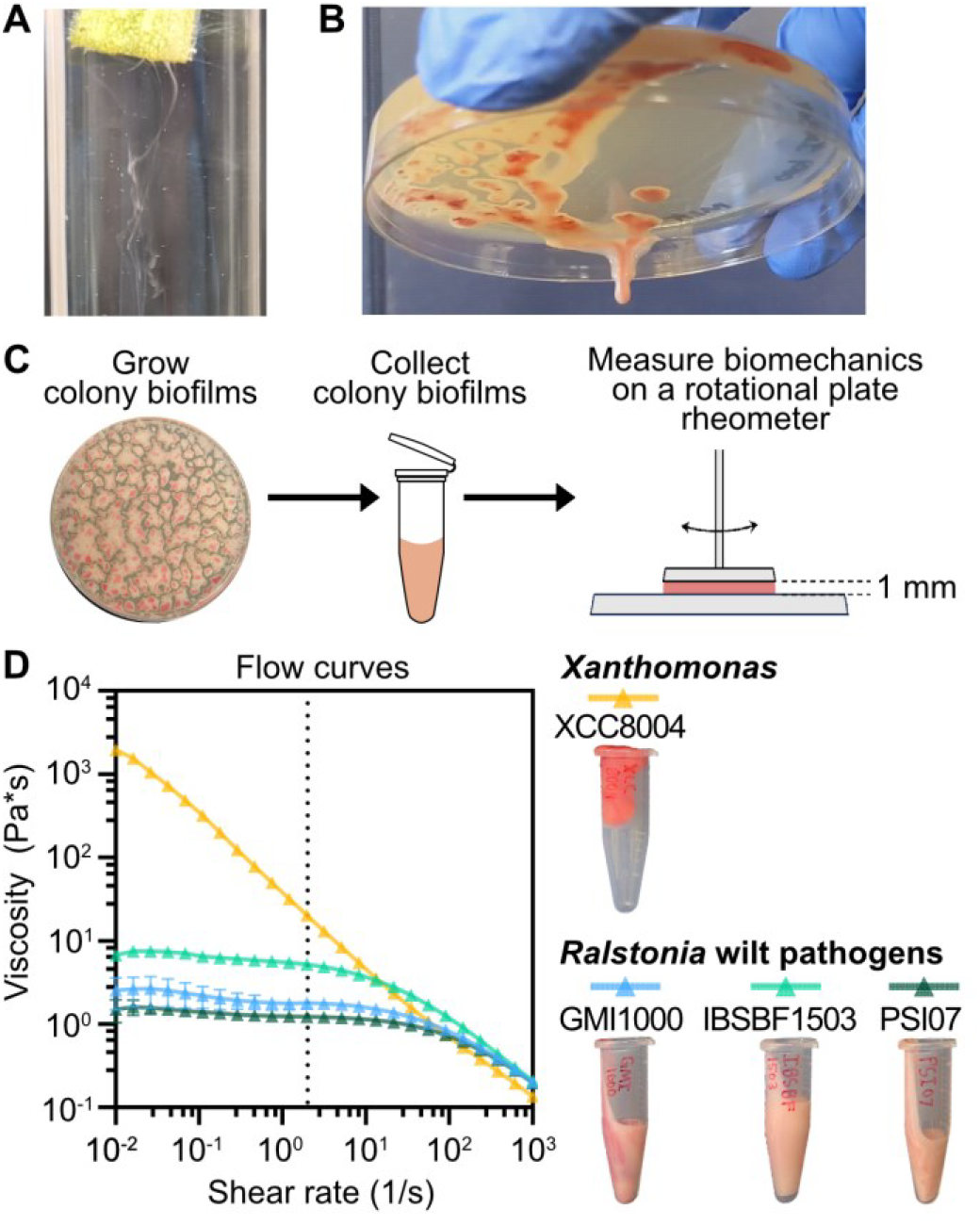
*Ralstonia solanacearum* species complex (RSSC) wilt pathogens produce biomass with viscous flow mechanical behavior. Photographs of (***A***) RSSC biomass (*Rpseu* GMI1000) streaming from a submerged, cut stem of a wilt symptomatic tomato plant and ***(B)*** RSSC colony biofilms (*Rpseu* GMI1000) that have coalesced and dripped upon inversion of petri dishes. The full video is available as **Video S1. *(C)*** Overview of biofilm rheology experiments. ***(D)*** Viscosity of colony biofilms of RSSC pathogens (*Rsol* IBSBF1503, *Rpseu* GMI1000, and *Rsyz* PSI07) and *Xanthomonas* XCC8004 at varying shear rates. The dashed line represents an estimated maximum shear rate within the xylem vessels of healthy plants (SI **“Results and Discussion”**) (45). Symbols show mean (N=3); error bars show SD.

**Fig. 2.**
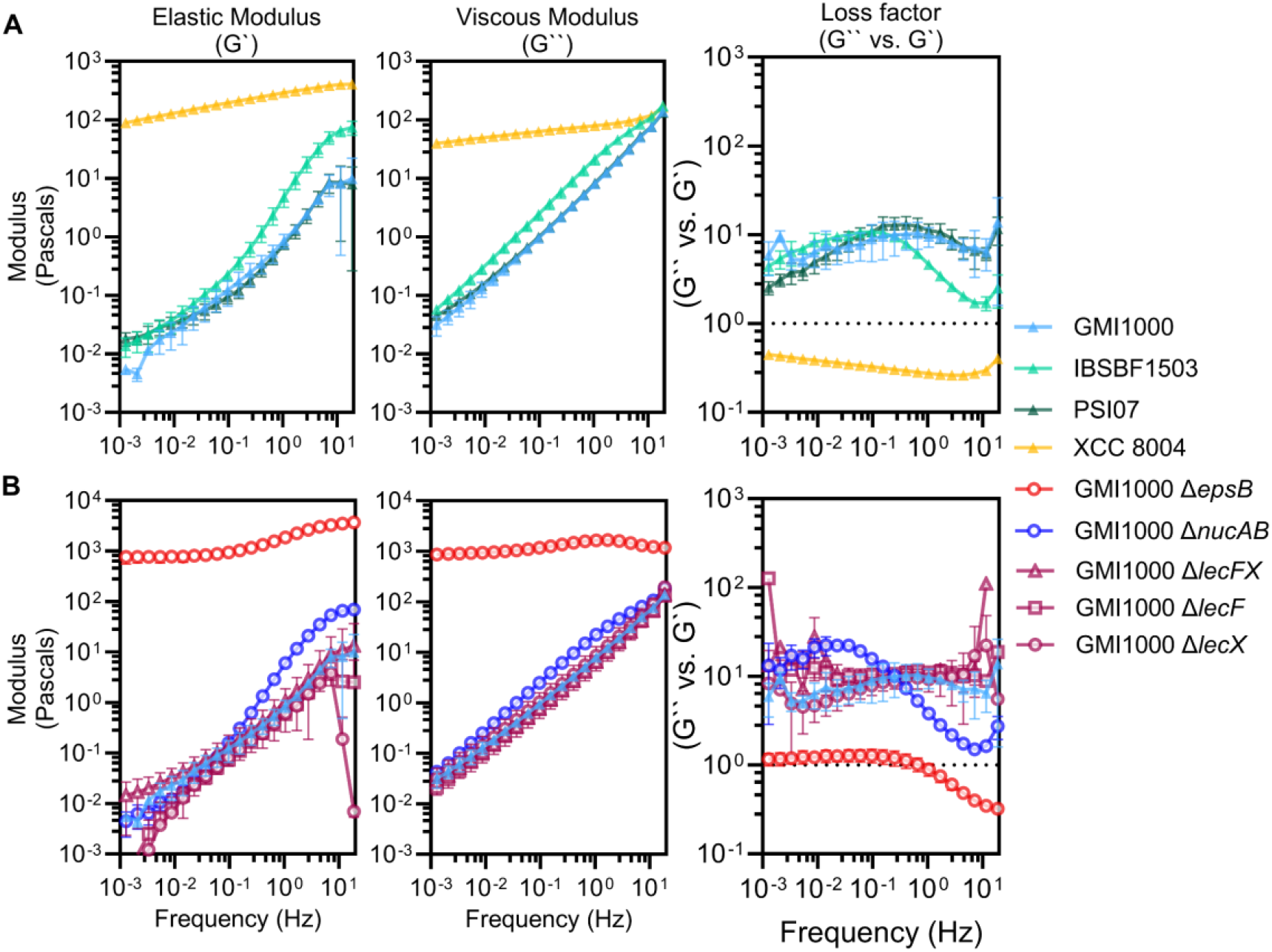
EPS-I confers unique viscoelastic mechanics to colony biofilms of RSSC wilt pathogens. Elastic (G`) and viscous (G``) moduli were measured in frequency sweep experiments and used to calculate the Loss factor for (***A***) wildtype RSSC wilt pathogens (GMI1000, IBSBF1503, and PSI07) and *Xanthomonas* (XCC8004) and (***B***) RSSC mutants with altered biofilm properties. Symbols show mean (N=3); error bars show SD. The dotted horizontal line represents the transition from elastic to viscous dominant behavior, corresponding to G`` / G`=1. Data from Wildtype GMI1000 is repeated across panel A and B plots for consistent comparisons

As a biologically informative comparison, we analyzed biofilms of a leaf-infecting *Xanthomonas* pathogen (*Xanthomonas campestris* pv. *campestris* 8004, “XCC8004”). Xanthomonads secrete xanthan gum, a model viscoelastic polymer in rheology (46, 47). As expected, the *Xanthomonas* biofilms were substantially more viscous than RSSC biofilms with a mean viscosity of 319 ± 3.22 Pa*s (± SD) at a shear rate of 0.11 s^-1^ (**Fig. 1D**).

Additionally, all wildtype RSSC and *Xanthomonas* biofilms were shear-thinning (**Fig. 1D**), meaning the viscosity decreased with increasing shear rate. Shear thinning is a typical trait for biofilms and can be contrasted with a shear thickening fluid such as a corn starch solution that resist deformation from strong forces, but readily flow when perturbed with weak forces (48). The shear-thinning feature of biofilms assists their ability to deform when external forces are high enough, flowing more easily with increasing flow rate in the xylem.

We next quantified the elastic (*G`*) and viscous (*G``*) moduli of RSSC and *Xanthomonas* biofilms in the linear viscoelastic region (see methods, SI “**Materials and Methods”** and **Fig. S1**, for details of rotational rheology). All strains exhibited a direct relationship between rotational frequency and the moduli, and *Xanthomonas* biofilms had higher elastic and viscous moduli than the RSSC biofilms at all frequencies (**Fig. 2A**). Moreover, the rate at which the viscous and elastic moduli increased as frequency increased was greater for RSSC biofilms than for *Xanthomonas* biofilms. For example, the elastic moduli of *Rsol* IBSBF1503 increased across frequencies about 5000-fold from a mean of 0.0137 Pa to 74.2 Pa. By contrast, *Xanthomonas* biofilms increased about 4.5-fold, from a mean of 89.1 Pa to 404 Pa. To compare the viscous and elastic moduli and characterize the biofilms as either viscoelastic solids or viscoelastic fluids, we determined the “loss factor” of the biofilms by calculating the ratio of G*``* to G*`*. Materials are elastic dominant (viscoelastic solids) if their loss factor is less than 1 and viscous dominant (viscoelastic fluids) if their loss factor is greater than 1. *Xanthomonas* biofilms were predominantly elastic at all driving frequencies (**Fig. 2A**). RSSC colony biofilms, on the other hand, were uniquely viscous-dominant with loss factors between 1.51 and 22.5, depending on the RSSC strain and measurement frequency (**Fig. 2A**). Overall, *Xanthomonas* biofilms are viscoelastic solids like all previously analyzed microbial biofilms (**Table S1**) (3–23). RSSC biofilms have viscoelastic fluid behavior where the mechanical work performed in deforming RSSC biofilms is lost as they flow rather than being stored and so they are unable to rebound like in an elastic material.

### The unique exopolysaccharide, EPS-I, is responsible for the viscous-dominant biomechanics of RSSC biofilms

The revelation that RSSC biofilms are viscoelastic fluids led us to ask, which matrix components confer these unique mechanics? We repeated the rheological assays with RSSC mutants that have altered biofilm composition: Δ*nucAB*, Δ*lecF*, Δ*lecX*, Δ*lecFX*, and Δ*epsB* mutants in the *Rpseu* GMI1000 background. The Δ*nucAB* mutant has elevated exDNA due to the absence of secreted DNases (29). The Δ*lecF*, Δ*lecX*, and Δ*lecFX* mutants lack lectin(s) that contribute structural stability to RSSC colonies (32). The Δ*epsB* mutant does not produce EPS-I because it lacks a Wzc family integral inner membrane protein that is required for EPS-I export (49). All mutant biofilms retained shear-thinning mechanical behavior (**Fig. S2**) (32). The DNase and lectin mutant biofilms exhibited moderately altered elastic (*G`*) and viscous (*G``*) moduli (**Fig. 2B**). Consistent with the Δ*nucAB* mutant’s development of dome-shaped colonies (29), Δ*nucAB* biofilm viscous moduli were up to 3-fold higher and elastic moduli up to 7-fold higher than wildtype GMI1000. In addition to their known increase in colony area and the reduction in biofilm viscosity (32), lectin mutant biofilms also had reduced elasticity. While these biofilm components clearly contribute to the viscoelastic behavior of RSSC biofilms, mutants lacking these components all still produced viscous dominant biofilms.

Notably, the Δ*epsB* mutant was the only mutant that lacked the characteristic viscous-dominant mechanics of wildtype RSSC biofilms with a loss factor between 0.308 and 1.53, depending on the measurement frequency (**Fig. 2B**). Additionally, the biofilms of the EPS-deficient mutant were dramatically more viscous and elastic than wildtype biofilms across all frequencies. At the median measured frequency (0.156 Hz), biofilm G` was 1,030 ± 199 Pa and 0.173 ± 0.0772 Pa for the Δ*epsB* mutant vs wild type, respectively (mean ± SD). Similarly, the biofilm G`` was 1,260 ± 91.0 Pa and 1.43 ± 0.177 Pa for the Δ*epsB* mutant vs wild type, respectively. These data reveal that EPS-I contributes to the viscous-dominant nature of RSSC biofilms, reducing the crossover strain required for RSSC biofilms to flow.

### EPS-I exopolysaccharide confers collective biofilm mobility of plant pathogenic RSSC biofilms in diverse environments

Our discovery that EPS-I confers unique rheological properties to RSSC biofilms motivated us to explore the mechanistic understanding of this major virulence factor because the precise pathogenic function of EPS-I remains cryptic. The **SI Results and Discussion** summarizes gaps in knowledge from prior *in planta* studies of EPS-I (24, 33, 42). Briefly, the literature was unclear about whether EPS-I contributes to pathogen growth *in planta*. Thus, we tested the contribution of EPS-I to pathogen growth and dissemination. Using cut petiole inoculations, we directly inoculated plant xylem with either wildtype GMI1000 or the Δ*epsB* mutant. After 3 days, population sizes were quantified at the site of inoculation and at distal sites 2-cm and 4-cm above and below the site of inoculation (**Fig. 3A**). At the site of inoculation, Δ*epsB* mutant had the same incidence as wild type (95%). However, at distal sites below and above the site of inoculation, there was lower incidence of the Δ*epsB* mutant: 69-81% wild type and 25-50% for the Δ*epsB* mutant. Overall, our results are consistent with a role of EPS-I as a dissemination factor.

**Fig. 3.**
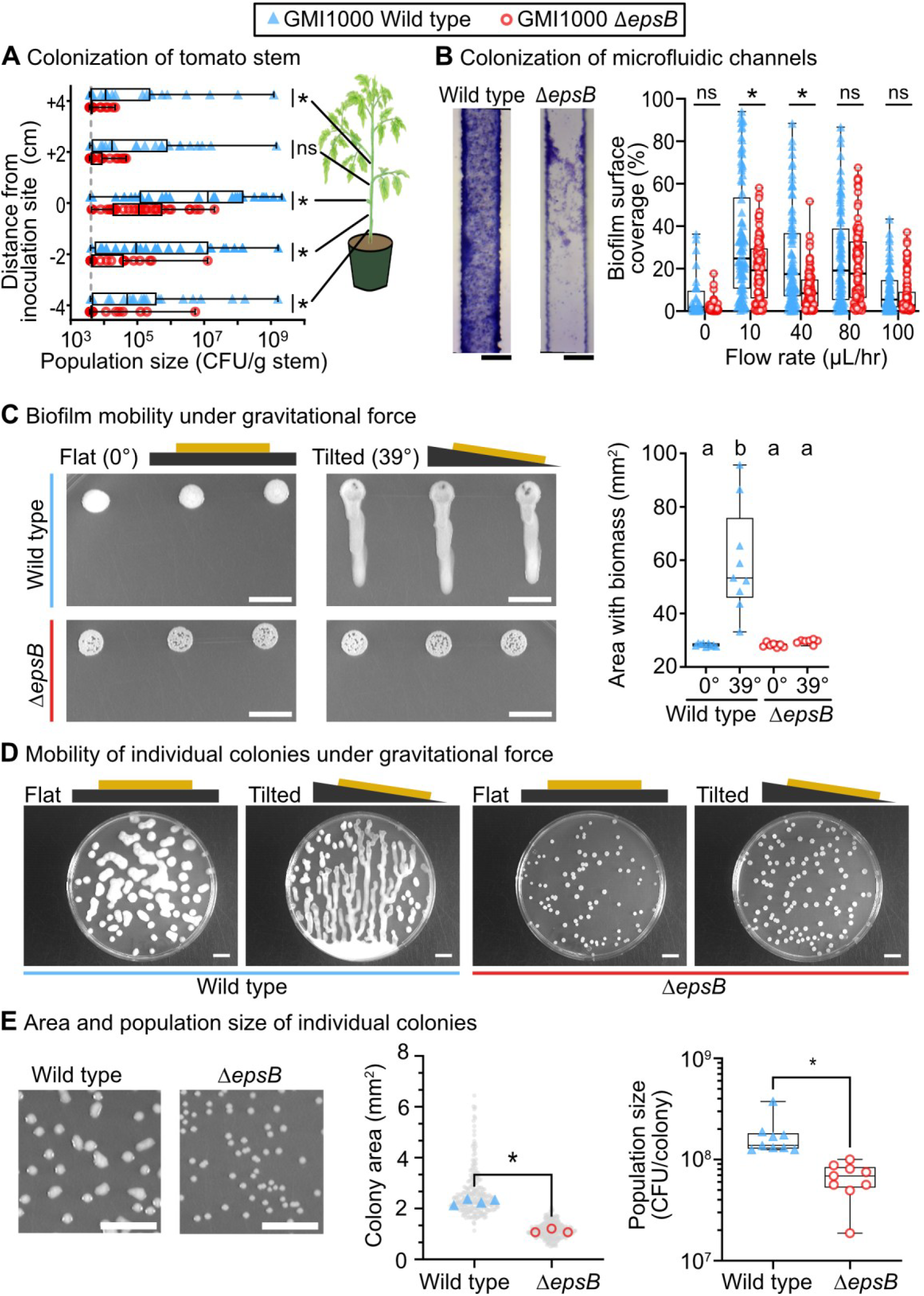
EPS-I confers biofilm mobility, which enhances RSSC fitness. ***(A)*** Dispersal of wildtype and Δ*epsB Rpseu* GMI1000 in tomato stems after cut-petiole inoculation (4 trials of n=8 plants per strain for a total of n=32 plants per strain; * represents *P* <0.05, Mann-Whitney test). ***(B)*** Biofilm occupancy of CMC-coated microfluidic channels by wildtype and Δ*epsB Rpseu* GMI1000, following 72 hr of rich media flowing at various rates. The image shows a representative result at 10 µL medium /hr from the middle portion of the microchannel, and images from all flow rates are shown in **Fig. S5** (Scale bar = 50 μm; * represents adjusted *P* <0.05, ANOVA with Šídák’s multiple comparisons test). ***(C)*** Biofilm mobility of wildtype and Δ*epsB Rpseu* GMI1000 when grown in static flat vs. tilted conditions. (Scale bar = 1 cm; letters indicate *P*<0.0001 by Welch’s ANOVA with Dunnett’s multiple comparisons) full data set in **Fig S6. *(D)*** Mobility of individual colonies grown rich agar. Colony biofilms grew at a normalized density of 125 CFU/spot for two days on either a flat or tilted angle. A full image set for all RSSC strains and *Xanthomonas* XCC8004 can be found in **Fig. S7. *(E)*** EPS-I enables colony expansion, which increases population sizes per colony (Scale bar: 1 cm; middle panel: shows a superplot of individual colonies (grey) and the median for each petri dish (blue or red symbols) where * indicates *P*<0.0001 by Welch’s ANOVA with Brown-Forsythe multiple comparisons; right panel: * indicates *P*<0.0001 by Mann-Whitney test).

Nevertheless, our experiments demonstrate that the Δ*epsB* mutant had a general growth defect at the site of inoculation (**Fig. 3A** and **Fig. S4**). Wildtype populations reached a geometric mean 4.8 x10^6^ CFU/g at the inoculation site, compared to Δ*epsB* population sizes of 2.0 x10^5^ CFU/g. This growth defect at the inoculation site demonstrates that movement is unlikely the sole virulence function of EPS-I. The growth defect suggests that biofilm viscoelasticity also protects the pathogens from the external chemical stresses imposed by the plant host. Thus, to directly test the contribution of EPS-I to movement of RSSC populations, we sought to investigate RSSC biofilm mobility in the absence of plant immune defenses.

Microfluidic devices simulate the highly confined hydrodynamic environment that RSSC pathogens experience in xylem vessels. To investigate the contribution of EPS-I to the colonization of xylem-like environments in a flow-dependent manner, we cultured wildtype GMI1000 and the Δ*epsB* mutant in microfluidic devices coated with carboxymethyl cellulose-dopamine (CMC) to further replicate the environment of xylem vessels (**Fig. S5**) (50). We found that total coverage of wildtype biofilms was significantly greater than Δ*epsB* biofilms at 10 and 40 µL/hr flow rates (**Fig. 3B**) but not significantly different at higher and lower flow rates.

Wildtype GMI1000 coverage of the microfluidic channels had a geometric mean of 20.3% and 12.3%, respectively. By comparison, the Δ*epsB* mutant covered the microfluidic channels to 10.7% and 5.04% for the respective flow rates. While we recently demonstrated that EPS-I was necessary for biofilm development in these microfluidic channels, our analysis of the rheological contributions of EPS-I led us to question whether the contribution of EPS-I depends on flow. Interestingly, the 10 and 40 µL/hr flow rates translate to dynamic processes in the 1-10 s^-1^ frequency range, where the Δ*epsB* mutant biofilm is elastic-dominant (**Fig. 2B**).

It is known that flow in microchannels creates shear forces that can cause biofilms to grow, restructure, or detach from walls. Even very small fluid shear stresses can initiate biofilm formation in some species by triggering the secretion of polysaccharides that aid biofilm formation (51, 52). Conversely, strong shear associated with high flow rates can suppress biofilm formation. Fast bulk flow can directly inhibit biofilm development by carrying cells at much faster rates than the cells can attach to the surface or indirectly by carrying nutrients away faster than the nutrients can diffuse to surface-attached cells. Thus, the effect of biofilm viscoelasticity on biofilm structure and behavior is most evident in intermediate, physiologically relevant shear stresses, which are approximately up to ∼10^-2^ Pa in the case of xylem flow (**SI “Results and Discussion”**). At these stresses, typical viscoelastic solid biofilms would undergo restructuring due to their solid-like mechanical behavior, which is characterized by very slow relaxation and can result in stiffening, softening, or yielding, depending on the species (51). Our findings that EPS-I contributes to colonization of microfluidic channels in a flow-dependent manner suggests that the unique viscous-dominant rheology of RSSC biofilms contributes to the enhanced surface coverage at physiologically relevant shear stresses.

Knowing that wildtype RSSC colony biofilms can drip when deformed by gravity (**Video S1**), we developed simple assays to compare mobility of RSSC biofilms. In one assay, cell suspensions were spotted on agar and colony biofilms developed while the plates were incubated at constant fixed angles of 0° (flat) or 39° (tilted). As expected, the Δ*epsB* mutant biofilms remained in place while gravity caused the wildtype biofilms on the tilted plates to deform and disseminate up to 2 cm (**Fig. 3C** and **Fig S6**). Thus, EPS-I production enables wildtype RSSC to flow as a viscoelastic fluid, a trait we call “biofilm mobility” hereafter. We repeated the tilt plate assay with plates inoculated at low cell density so that individual cells would grow into spatially separated colonies. Once again, wildtype biofilms were mobile and dripped when grown on tilted agar while the Δ*epsB* biofilms remained in place (**Fig. 3D** and **Fig. S7**). Only the Δ*epsB* mutant and *Xanthomonas* XCC8004 biofilms lacked mobility (**Fig. 3D** and **Fig. S7**). Intriguingly, the images suggest that when multiple wildtype RSSC biofilms converged, the collective mass traveled longer distances as a confluence. This visualization of RSSC biofilm mobility underscores the demonstrated fact that EPS-I production is a collective good that is produced in a quorum sensing-dependent manner (38–40).

### Biofilm mobility improves RSSC fitness by enabling the pathogen to passively expand its niche

Although EPS-I production is a net benefit to RSSC fitness *in planta* (**Fig. 3A**), this outcome is context-dependent. When grown in liquid media, EPS-I production reduces RSSC’s maximum growth rate (**Fig. S8**) (53), almost certainly due to the metabolic burden of synthesizing and exporting the EPS-I polymer. We hypothesized that EPS-I is a net benefit to RSSC fitness in conditions where the viscoelasticity allows RSSC biofilms to expand their physical niches and gain access to more nutrients. We tested this hypothesis on agar plates, quantifying colony area and population sizes. Even when growing flat, gravity deforms wildtype biofilms, causing colony expansion (**Fig. 3E**). We used CellProfiler to quantify the area of individual colonies (54), which demonstrated that wildtype RSSC colonies occupy roughly double the area compared to Δ*epsB* colonies (**Fig. 3E**). Wildtype GMI1000 colonies grew to a geometric mean of 2.48 mm^2^, and the Δ*epsB* colonies grew to 1.11 mm^2^. However, a larger colony does not necessarily mean that wildtype cells had higher reproduction. In addition to lacking EPS-I, Δ*epsB* colony biofilms had a significant decrease in moisture content (**Fig. S9**). Wildtype GMI1000 colonies had a moisture content of 90.3% ± 0.251% (mean ± SD), while Δ*epsB* colony biofilms had a mean moisture content of 64.6% ± 0.286%, suggesting the difference in colony size might be confounded to a difference in water. Thus, we directly measured the population size of individual colonies by excising colonies with a blade, homogenizing the colony, and dilution plating to enumerate colony forming units (CFUs). This direct test of bacterial fitness demonstrated that individual wildtype colonies contained approximately 4-fold more cells than Δ*epsB* colonies (**Fig. 3E**). Wildtype GMI1000 colony populations contained a geometric mean of 1.6x10^8^ CFU/colony, whereas the Δ*epsB* colonies contained 6.0x10^7^ CFU/colony.

We assume that EPS-I production had the same metabolic cost whether the cells grow in liquid broth or on agar. Nevertheless, wildtype cells on agar were able to reproduce more than the Δ*epsB* mutants. Why? Although we cannot rule out the hypothesis that the Δ*epsB* cells were constrained by water stress in the absence of hygroscopic EPS-I, we speculate that production of EPS-I allowed the colonies to increase the cells’ access to nutrients via a microstructural change closely linked to biofilm rheology. The resulting flowability increased the area of a colony as it spreads due to gravity, enhancing nutrient flux across the agar-colony interface. Additionally, we speculate that the elastic modulus (G`) roughly indicates the amount of thermal energy stored per unit volume of biofilm matrix. The relevant volume in this case is set by the typical mesh size within the scaffolded structure of the entangled extracellular polymer matrix (55). G` is then inversely related to the volume of the space between polymeric entanglements. A reduction in the elastic modulus due to EPS-I then indicates an increase in the volume available for nutrients to reside in and diffuse freely into without the close confinements of a tight polymeric mesh. Thus, the increased opportunity to acquire nutrients exceeds the cost of EPS-I production.

### Viscoelastic fluid biofilms are a key evolutionary event that coincides with the emergence of wilt pathogenesis within the *Ralstonia* genus

RSSC wilt pathogens are a monophyletic lineage within the *Ralstonia* genus. Other species of *Ralstonia* are found in habitats such as soil, the plant rhizosphere, surface water, industrialized water systems such as the water recycling system on the International Space Station, and opportunistic infections of humans. To explore the evolutionary history of the RSSC wilt pathogens’ EPS-I dependent, viscoelastic liquid biofilms, we explored publicly available genomic data and biofilm mobility phenotypes of diverse isolates across the *Ralstonia* genus.

RSSC isolates produce and export EPS-I via the products of the 17-gene *eps* cluster that is encoded on a megaplasmid (49, 56, 57). We used BLASTp to search 399 RSSC genomes and 72 non-RSSC *Ralstonia* genomes for homologs of the *eps* cluster genes and the neighboring *xpsR* gene whose product activates expression of the *eps* genes (38). Within the plant pathogenic RSSC, the *eps* genes are syntenic and nearly universally conserved (**Fig. 4A-B** and **Fig S10**), except for a single genome (**Fig. S11)**. Most genes had high amino acid sequence identity with a minimum identity of 84.5%-96.5% to the query, but RSp1005 had higher variation with identity as low as 65.1%. None of the non-RSSC *Ralstonia* genomes encoded the *eps* gene cluster although low identity homologs were occasionally detected (**Fig. 4B**). A similar analysis suggests that the *nucA* nuclease is also specifically present in RSSC genomes while the *nucB* nuclease is conserved across the entire genus (**Fig. S12**).

**Fig. 4.**
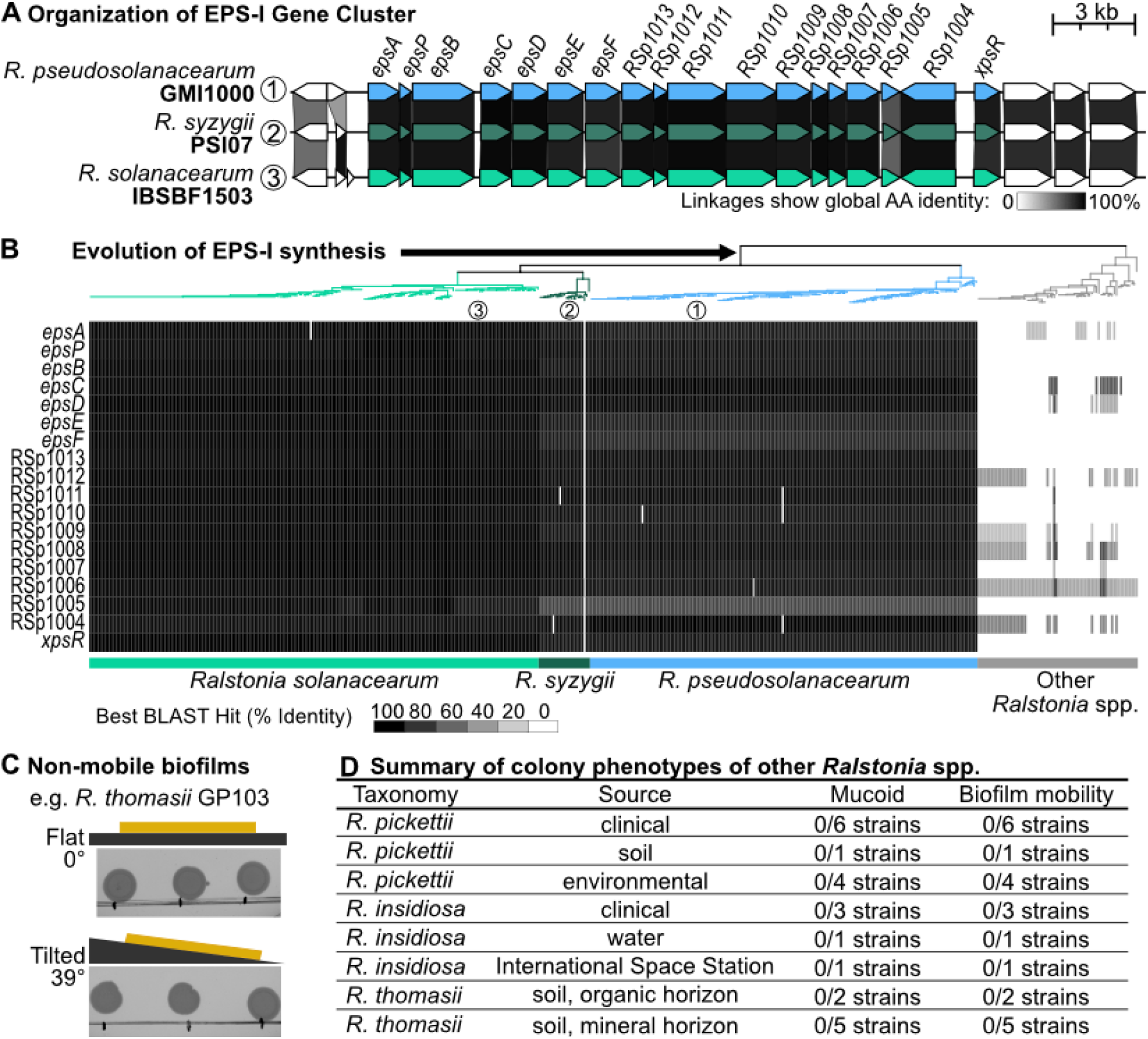
EPS-I production and biofilm mobility is an evolutionary innovation of RSSC wilt pathogens. ***(A)*** The *eps* gene cluster synteny of strains from the three RSSC species was visualized with Clinker. ***(B)*** Presence of *eps* genes in genomes of the RSSC pathogens and non-RSSC *Ralstonia* species that do not wilt plants. Rectangles show percent amino acid identity of the best BLASTp hit for queries with *Rsol* K60 proteins. The approximate maximum likelihood phylogenetic tree was built from 49 conserved bacterial COGs using the KBase Insert Genomes into SpeciesTree v2.2.0 application and visualized in iToL. ***(C-D)*** Results of biofilm mobility assay non-wilt pathogenic *Ralstonia*. Full results are in **Fig. S15** (***C***) Representative results showing the soil isolate *R. thomasii* GP103 ***(D)*** Summary of colony biofilm phenotypes of non-wilt-pathogenic *Ralstonia* isolates.

The pattern of *eps* gene presence suggests that either 1.) the *eps* genes were acquired or evolved in the common ancestor of the RSSC when this lineage diverged from the rest of the genus, or 2.) the ancestor of the entire *Ralstonia* genus encoded the *eps* gene cluster, and the non-RSSC lineage lost the *eps* genes. To investigate these evolutionary hypotheses, we used a cluster reconstruction and phylogenetic analysis pipeline (58, 59) to reconstruct phylogenies for the 17 RSSC *eps* cluster genes that encode the proteins for synthesis and export of EPS-I. Each EPS protein was BLASTp searched against a database of 16,355 betaproteobacterial genomes, including 508 RSSC genomes, and phylogenies of each gene were constructed after the BLAST datasets were truncated through sequence similarity clustering (58). This analysis identified 1,563 non-RSSC betaproteobacterial genomes that had 2 or more *eps* genes within 20 kb, but only the RSSC pathogen genomes encoded the full *eps* gene cluster. Several *eps* genes were rarely detected in the betaproteobacterial genomes: *epsE, epsF*, RSp1013, RSp1015, and RSp1003 (**Fig. S13**). Several non-RSSC genomes had as many as 13 homologs to the 17 *eps* genes (**Fig. S13**), but those gene clusters contained additional non-homologous genes and often varied in gene organization (**Fig. S14**), suggesting the gene products would produce a different exopolysaccharide than EPS-I. The gene phylogenies provided a second line of evidence that the *eps* cluster is conserved across the RSSC and shares a recent origin within the clade of RSSC wilt pathogens. The outgroups of the RSSC *eps* genes were commonly identified in *Cupriavidus*, the sister genus of *Ralstonia*. There was also evidence of recombination of partial *eps*-like clusters in sporadic genomes of *R. pickettii* and *R. mannitolilytica* and throughout the Alcaligenaceae and Burkholderiaceae. As a third approach, we used cblaster to query the 17 *eps* genes against public genomes in NCBI’s nr database. Once again, the *eps* gene cluster was only identified in RSSC genomes. Overall, the gene cluster responsible for EPS-I production is unique to the RSSC. This result aligns with the effectiveness of antibody-based diagnostics to accurately detect bacterial wilt disease through the identification of EPS-I (60) .

Together, the evolutionary genomic analyses strongly suggest that production of viscous-dominant EPS-I is unique to the RSSC. But what about the physiology? To determine whether the non-RSSC *Ralstonia* shared the trait of forming viscoelastic fluid colony biofilms, we investigated the biofilm mobility phenotype of other *Ralstonia* spp. isolated from the International Space Station, soil, and human patients. All 23 of these isolates lacked biofilm mobility (**Fig. 4C-D** and **Fig. S15**), which demonstrates that the mobile, fluid biofilm trait is specific to the RSSC wilt pathogens.

### An improved model of the role of EPS-I in RSSC virulence

We propose that EPS-I is a key *in planta* movement factor that transforms RSSC biofilms into viscoelastic fluids. Based on the measured viscosities of wildtype RSSC biofilms (**Fig. 1D**) and estimated shear rates in xylem vessels (45), we calculate that viscous flow could passively disseminate RSSC biofilms in the range of centimeters per day while the Δ*epsB* mutant is practically immobile against flow (**SI “Results and Discussion”**). The chemical structure of EPS-I sheds light on the unique mechanics. Typical polysaccharides are composed of polar sugars. However, EPS-I is an amphiphilic polysaccharide composed of trimeric repeats of N-acylated derivatives of galactosamine, deoxygalacturonic acid, and the reduced sugar bacillosamine (**Fig. S16**) (31). We suspect that when in the extracellular matrix, this polymer has minimal inter- and intra-molecular hydrogen bonding when compared to canonical polysaccharides, leading to a decrease in biofilm elasticity and viscosity. As an amphiphilic polymer, EPS-I appears to play an analogous role in bacterial movement as rhamnolipids and other small molecule biosurfactants that coat surfaces and enable cells to slide (61). Our results suggest that EPS-I is analogous to a biosurfactant that enables biofilms to expand by flow when deformed by the modest shear forces within xylem vessels.

Thus, RSSC pathogens have three movement mechanisms: flagellar swimming motility, type IV pilus-driven twitching motility, and EPS-I dependent biofilm mobility. RSSC predominantly use swimming motility to invade roots (62). Swimming motility is repressed in the xylem (62), and non-flagellated mutants have wildtype virulence and normal *in planta* dissemination (62, 63). Multiple lines of evidence suggest that pilus-mediated motility and EPS-mediated biofilm mobility have complimentary roles in RSSC dissemination through plant hosts. Even when direct inoculation into plant stems allows the pathogen to bypass root invasion, type IV pili and EPS-I are both strongly required for virulence (33, 64) and dissemination (**Fig. 3**) (42, 63).

The evolutionary conservation of EPS-I biosynthesis across the three RSSC species suggests that biofilm fluidity contributes to these pathogens’ success. Biofilm mobility has also been demonstrated for the distantly-related xylem pathogen *Erwinia amylovora* although the rheological properties for biofilms of this pathogen have not been analyzed (65). Not all vascular plant pathogens have mobile biofilms. Here, we demonstrated that the vascular black rot pathogen *Xanthomonas* XCC8004 produces highly elastic, viscoelastic solid biofilms, similar to all previously characterized bacterial and fungal biofilms. These *Xanthomonas* biofilms were non-mobile under the force of gravity. However, a key principle of rheology is “everything flows;” with sufficient strain, viscoelastic solid biofilms will migrate by viscous flow (2). Nevertheless, RSSC pathogens can fully wilt a plant host within days, which is dramatically faster than *Xanthomonas* and all other bacterial or fungal wilt pathogens. Thus, viscoelastic fluid biofilms are key factors for systemic *in planta* spread of RSSC pathogens.

### Materials and Methods

Detailed methods for bacterial strains and culture media and experiments on rheometry of colony biofilms, bacterial growth and dispersal in tomato stems, biofilms in CMC-coated microfluidic, biofilm mobility on agar, image analysis with CellProfiler, growth curves, and bioinformatic analyses are present in **SI Appendix, Supplementary Methods**.

## Supporting information

Video S1 - Real time video of RSSC biofilm mobility from Agar

## Acknowledgments

We thank Prof. Cooper Battle (Willamette University) and Clay Fuqua (Univ. Indiana) for valuable discussion. We also thank Prof. Pamela Ronald (University of California, Davis) for the contribution of the *Xanthomonas campestris* pv. campestris 8004 isolate.

## Data availability

All data in this publication can be found on Zenodo (https://zenodo.org/) with the DOI: 10.5281/zenodo.14767251

## Author Contributions

MCA: Conceptualization, Analysis, Experiments / Investigation, Methodology, Visualization, Writing – original draft, Writing – review & editing; JL: Conceptualization, Analysis, Experiments / Investigation, Methodology, Visualization, Writing – original draft, Writing – review & editing; TML: Conceptualization, Data curation, Analysis, Funding acquisition, Methodology, Resources, Supervision, Visualization, Writing – original draft, Writing – review & editing; HM: Conceptualization, Analysis, Funding acquisition, Methodology, Resources, Supervision, Writing – original draft, Writing – review & editing; NA: Analysis, Experiments / Investigation, Methodology, Visualization, Writing – review & editing; TMT: Conceptualization, Data curation, Analysis, Funding acquisition, Experiments / Investigation, Methodology, Resources, Supervision, Validation, Writing – review & editing; TC: Analysis, Experiments / Investigation, Visualization, Writing – review & editing; MDC: Conceptualization, Resources, Writing – review & editing; CA: Conceptualization, Resources, Supervision, Writing – review & editing; ZK: Data curation, Analysis, Experiments / Investigation, Methodology, Validation, Visualization, Writing – review & editing; JJ: Funding acquisition, Supervision, Writing – review & editing; AC: Experiments / Investigation, Resources, Visualization, Writing – review & editing; KMD: Funding acquisition, Resources, Supervision, Writing – review & editing; SW: Experiments / Investigation, Writing – review & editing; MJW: Experiments / Investigation, Resources, Writing review & editing; LJC: Resources, Writing – review & editing; NW: Data curation, Analysis, Experiments / Investigation, Methodology, Validation, Visualization, Writing – review & editing; LB: Experiments / Investigation, Writing – review & editing; LTC: Experiments / Investigation, Writing – review & editing

## Funding Information

None of the funders had any role in study design, data collection and analysis, decision to publish, or preparation of the manuscript.

| **Source** | **Grant Reference Number** | **Authors** |
| --- | --- | --- |
| USDA Hatch | #1023861 | Tiffany Lowe-Power |
| USDA Hatch Multistate | #7004345 | Kristen M. DeAngelis |
| UC Davis Academic Senate |  | Tiffany Lowe-Power<br>Hari Manikantan |
| USDA NIFA | #2024-67011-42914 | Matt Cope-Arguello |
| USDA NIFA | #2022-67013-36272 | Tiffany Lowe-Power |
| U. of South Alabama<br>Office of Research &<br>Economic Development | #A24-0097-001 | Tuan M. Tran |
| USDA Specialty Crop<br>Block Grant | #23SCBGPAL 1173-00 | Tuan M. Tran |
| USDA NIFA | #2024-67012-42635 | Nathalie Aoun |
| USDA Hatch | #7004345 | Kristen M. DeAngelis |
| Cystic Fibrosis Foundation<br>Funding | CAVERL22AB0 | Lindsay J. Caverly |
| NSF Graduate Research<br>Fellowship | DGE-1747503 | Mariama Carter |

