## Supporting Information for "The EPS-I exopolysaccharide transforms *Ralstonia* wilt pathogen biofilms into viscoelastic fluids for rapid dissemination *in planta*"


Author order:

- [1]Matthew Cope-Arguello
- [2]Jiayu Li
- [3]Zachary Konkel
- [1]Nathalie Aoun
- [1]Tabitha Cowell
- [5]Nicholas Wagner
- [6]A. Li Han Chan
- [7]Lan Thanh Chu
- [2]Samantha Wang
- [8,9]Mariama D. Carter
- [8]Caitilyn Allen
- [10]Lindsay J. Caverly
- [7]Loan Bui
- [6]Kristen DeAngelis
- [11]Matthew J. Wargo
- [5]Tuan M. Tran
- [3,4]Jonathan M. Jacobs
- [2]Harishankar Manikantan*
- [1]Tiffany M. Lowe-Power*

| |
|---|
| [1]Department of Plant Pathology, University of California, Davis, Davis, CA, USA 95616 |
| [2]Department of Chemical Engineering, University of California, Davis, Davis, CA, USA 95616 |
| [3]Infectious Diseases Institute, The Ohio State University, Columbus, OH, 43210, USA |
| [4]Department of Plant Pathology, The Ohio State University, Columbus, OH, USA |
| [5]Department of Biology, University of South Alabama, Mobile, AL 36688 |
| [6]Department of Microbiology, University of Massachusetts, Amherst, MA, USA 01003 |
| [7]Department of Biology, University of Dayton, Dayton, OH 45469 USA |
| [8]Department of Plant Pathology, University of Wisconsin-Madison, Madison, WI 53706, U.S.A. |
| [9]Department of Crop Genetics, John Innes Centre, Norwich Research Park, Norwich NR4 7UH, United Kingdom |
| [10]Department of Pediatrics, University of Michigan Medical School, Ann Arbor, MI 48109 |
| [11]Department of Microbiology & Molecular Genetics, University of Vermont Larner College of Medicine, Burlington, VT 05405 |

* Harishankar Manikantan and Tiffany M. Lowe-Power (co-corresponding authors)

### Supporting Information Text

### SI Results and Discussion

#### Dissecting the role of EPS-I in RSSC growth vs. dissemination *in planta.*

Several papers and reviews claim that EPS-I is dispensable for RSSC growth *in planta* (1–3). However, our results and careful dissection of the published results contradict this conclusion. Here, we explain our re-analysis of this literature. A review states, "There is now general agreement that, despite [*eps* mutants] multiplying normally in stems near the site of wound inoculation" (1). That study's results state "The marked reduction in virulence of some EPS-deficient strains was not due to a failure to multiply within stems of tomato plants," citing a figure that shows an XY line graph of wildtype vs. an *eps* mutant population size over time, after a stem inoculation, where the population size is reported as "mean" "CFU per 9-cm stem segment centered on the site of inoculation" (2). Although the reported means are nearly identical at each time point, it is not clear whether the CFU data was properly log-normalized before averaged. The same research group revisited the population growth of *eps* and wildtype strains in Saile *et al*. 1997, which states in the abstract "[*eps*] mutants multiplied as well as wild type *in planta*" while the results demonstrate and state the contradictory finding "[the *eps* mutant] AW1-1 multiplied less than the wild type [at the site of inoculation]" (3). Although this result is difficult to interpret due to the visualization of the data as a 3D bar chart, the population sizes at the site of inoculation for wild type appear to be 3 plants at ~$10^{11}$ CFU/g, 1 plant at ~$10^{10}$ CFU/g, and 1 plant at ~$10^4$ CFU/g, whereas population sizes for the *eps* mutant appear to be 2 plants at ~$10^9$ CFU/g, 1 plant at ~$10^7$ CFU/g, and 2 plants at ~$10^5$ CFU/g.

Thus, we robustly interrogated this question by analyzing population size and spread of wildtype GMI1000 and the Δ*epsB* mutant across 4 trials of 8 plants per strain per trial (**Fig. 3A**). In each of our trials, the Δ*epsB* mutant had lower median and geometric mean population sizes than wild type at the site of inoculation (Summary statistics of population sizes [CFU/g], per trial: wildtype geometric means: $6.7\times10^8$, $9.1\times10^8$, $3.1\times10^8$, $5.0\times10^9$; Δ*epsB* geometric means: $1.4\times10^7$, $8.6\times10^6$, $1.0\times10^7$, $2.1\times10^7$; wild type medians: $6.6\times10^9$, $4.8\times10^9$, $1.0\times10^9$, $6.7\times10^7$; Δ*epsB* medians: $2.5\times10^7$, $3.2\times10^6$, $4.2\times10^6$, $2.5\times10^7$). If the Δ*epsB* mutant and wild type truly had equal growth in stems, we could directly attribute the Δ*epsB* mutant's lower population sizes at distant locations (2 and 4 cm above/below the inoculation) to a defect in the ability of the Δ*epsB* mutant to spread. However, the Δ*epsB* mutant evidently has a growth defect *in planta,* which means it is difficult to directly tie EPS-I production to RSSC movement *in planta.*

#### Estimating shear rates and stresses in xylem tubes and microfluidic channels.

Flow of sap within xylem-like tubes is well approximated by the Hagen-Poiseuille equation (4–6) which relates liquid flow rate ($Q$ ) linearly to the pressure drop across the length of the vessel ($\Delta P/L$ ), inversely to fluid viscosity ($\mu$), and to the fourth power of tube radius $R$:

$$Q = \frac{\Delta P}{L}\frac{\pi R^4}{8\mu}. \tag{1}$$

A laminar flow within such a tube can be solved using standard techniques of fluid dynamics to give the fluid velocity through the channel as a function of radial location ($r$ ) within the tube:

$$u(r) \ = \ \frac{\Delta P}{L}\frac{1}{4\mu}[R^2 - r^2], \tag{2}$$

If the entire volume of the tube is filled with the biofilm material while still being driven by the same pressure gradient, the resulting flow rate becomes

$$Q_b = \frac{\mu}{\mu_b}Q, \tag{3}$$

where the subscript $b$ denotes biofilm viscosity or flow rate, and the quantities without subscript are those of the sap. At low shear rates, the viscosity of RSSC biofilm is ~1 Pa.s (**Fig. 1D**), which is about a 1000-fold higher than that of clean sap [~$10^{-3}$ Pa.s (6)]. The resulting flow rate $Q_b$ within a biofilm-filled tube is then smaller by a factor of 1000 compared to that of pure sap.

Typical clean sap flow rates are in the range of $10^{-15}$ -$10^{-13}$ $m^3/s$ (4, 7). The same xylem or microfluidic channels filled with RSSC biofilm and driven by the same pressure gradient will then result in biofilm flow rates in the range of $10^{-18}$ -$10^{-16}$ $m^3/s$. The maximum velocity in such a flow can be estimated from Eq. (2) and using Eq. (1) to substitute for the pressure gradient as

$$u_b = \frac{2Q_b}{\pi R^2}. \tag{4}$$

Substituting estimated biofilm flow rates and using a typical channel radius of $10^{-5}$ m gives biofilm flow velocity in the range of $10^{-8}$ -$10^{-6}$ m/s. This translates to millimeters to centimeters per day of passive spread due to flow, consistent with observed motility in **Fig. 3A**.

We have so far assumed that the biofilm material occupies the entire channel, which might not be the case in early stages of infection. If an adherent biofilm coats the channel walls to a small thickness $\delta$, the effective radius of the available channel decreases from $R$ to $R - \delta$. The volume of the channel available for sap to flow decreases and the smaller radius suppresses the maximal fluid velocity as well as the flow rate.

A more conservative estimate for very thin biofilms that adhere to the channel walls can be determined by evaluating the viscous stress exerted by the predominantly sap-filled flow in the channels. Using data from Bouda *et al.* 2019 (4), we estimate strain rates ($\dot{\gamma}$) in the range of 0.1 to 10 1/s in xylem tubes. For microfluidic measurements, we may still use the same expression for an order of magnitude estimate despite the channels being square in cross-section. For example, at 10 μL/hr across 20 parallel channels of 20 μm-by-20 μm cross section, we obtain a strain rate in the range of 10 1/s near the walls, representing the higher end of xylem relevant shear flows. The associated viscous stress is then

$$\sigma = \mu\dot{\gamma} = \frac{4Q\mu}{\pi R^3}, \tag{5}$$

which is up to ~$10^{-2}$ Pa in xylem. Since the biofilm is much more viscous than xylem sap (~1 Pa.s vs. ~$10^{-3}$ Pa.s at low shear rates) and if the biofilm layer is thin relative to the tube diameter, we can assume that the bulk flow and resulting wall shear stress exerted by the sap is comparable to those in a clean channel. Such a shear stress, when exerted at the sap-biofilm interface, generates a strain rate within the biofilm given by

$$\dot{\gamma} = \frac{\sigma}{\mu_b}, \tag{6}$$

assuming the viscous behavior of the biofilm follows a generalized Newtonian behavior. For slow flows, using a plateau viscosity of ~1 Pa.s for wildtype RSSC (**Fig. 1D**), this gives a strain rate of the order of $10^{-2}$ 1/s. For wall-adherent biofilms that are micrometers thick within microfluidic channels or xylem tubes, this translates to a velocity of about $10^{-2}$μm/s or a few millimeters a day. For the same pressure drop, biofilms that fill up the vessel will be driven faster than when it merely coats the walls because all the stress due to the applied pressure acts to move the biofilm rather than primarily move the sap which then tugs on a thin viscous biofilm layer at the walls.

These estimates represent a lower limit of spread due to fluidity of the biofilm. The biofilm's effective viscosity will likely drop upon imbibing sap, which will result in a faster flow rate following Eq. (3), and faster colony motility. Further, the thickness of the biofilm increases over time until the channel is mostly occupied by biofilm and then Eq. (4) is a better estimate of the speed of spread.

In summary, wildtype RSSC viscosities measured here support the improved model of biofilm motility we present in the main text. RSSC biofilms lacking EPS have viscosities that are a factor of 100 or more relative to the wild type, and so the flow rate and resulting viscous spread is smaller by the same factor. In other words, the Δ*epsB* mutant biofilm and the comparably viscous *Xanthomonas* biofilm are practically immobile if motility is driven by viscous flow alone.

### Materials and Methods

#### Bacterial Strains and Media

Bacteriological media included rich CPG broth (1 g/L casamino acids, 10 g/L bacto-peptone, 5 g/L glucose or dextrose, and 1 g/L yeast extract), CPG+TZC agar (CPG broth with 15 g/L agar and 20 mg/L tetrazolium chloride) (10), the original Boucher's minimal medium (BMM; 0.5 g/L ammonium sulfate, 50 mg/L magnesium sulfate heptahydrate, 0.125 mg/L iron (II) sulfate heptahydrate, 3.4 g/L potassium phosphate monobasic, pH adjusted to 6.5 with 4 M potassium hydroxide) (11), or improved BMM (iBMM, which is BMM supplemented with micronutrients: 5 mg/L iron (II) sulfate heptahydrate, 50 mg/L EDTA disodium salt dihydrate, 22 mg/L zinc sulfate heptahydrate, 11.4 mg/L boric acid, 50.4 mg/L manganese (II) chloride tetrahydrate, 1.6 mg/L cobalt (II) chloride hexahydrate, 1.56 mg/L copper (II) sulfate pentahydrate, 1.12 mg/L ammonium heptamolybdate tetrahydrate) (12). All culture media were prepared using Milli-Q-filtered, deionized water as a base. Minimal media were filter-sterilized using filters with 0.22 μm pores and CPG medium was autoclaved.

Bacterial strains used in this study are listed in **Table S2**. All mutants were in the *R. pseu* GMI1000 background and consist of Δ*epsB* (RS_RS22035), Δ*nucAB* (lacking *nucA* RS_RS03750 and *nucB* RS_RS12310), Δ*lecF* (RS_RS10890), Δ*lecX* (RS_RS26140), and Δ*lecFX* (11, 12). Strains were cryopreserved as a cell suspension in 37.5% glycerol at -80 °C. Cell density of bacterial suspensions was approximated by measuring the absorbance of 600 nm light ($A_{600}$) with an

Ultrospec 10 Cell Density Meter (Amersham Biosciences). Density adjustments were based on a conversion where $A_{600}$ = 1 corresponds to a cell density of $5x10^8$ CFU/mL for all *Ralstonia* strains and a cell density of $1.6x10^9$ CFU/mL for *Xanthomonas* XCC8004. To adjust cell densities, suspensions were diluted with Milli-Q filtered water, except for *Xanthomonas campestris* XCC8004, which was diluted using the original BMM recipe without added carbon sources.

**Colony Biofilm Rheometry Experiments**
To culture colony biofilms, strains were streaked for isolated colonies from glycerol stocks onto a CPG+TZC agar plate and incubated at 28 °C for two days or at room temperature (~22 °C) for three days. For each replicate, an individual colony was inoculated into CPG broth and incubated at 28 °C with 250 rpm shaking overnight. These cultures were used to grow 5000 CFU per agar plate. To do so, cell density of the cultures was adjusted to 5 x $10^4$ CFU/mL, 100 µL of the cell suspension was spread onto a minimum of four CPG+TZC agar plates using sterile glass beads, and plates were incubated for three days at 28 °C. Colony biofilms were harvested from the replicate plates into 1.7 mL microcentrifuge tubes using a glass rod. A minimum of 1 mL of colony biofilm was collected for each sample. To minimize moisture loss, tubes were sealed whenever possible.

Mechanical properties of colony biofilm samples were measured using an Anton Paar Modular Compact Rheometer 302, a 25 mm parallel plate (PP 25) measuring system, and RheoCompass software. For all assays, the rheometer base plate was set to 28°C. Approximately 1 mL of sample was pipetted onto the base plate. The plate was lowered to a gap of 1.05 mm, excess biomass was trimmed, and the plate was lowered to a final gap of 1 mm.

Flow curve experiments were performed to determine the viscosities of the colony biofilms. During a flow curve, the measuring system rotates in a consistent direction at increasing shear rates. For each flow curve experiment, 25 data points were collected as the shear rate increased logarithmically from 0.01 - 1000 $s^{-1}$. Viscosity data were recorded when viscosity reached equilibrium for each shear rate, with a maximum measuring period of 30 s. Three separately prepared biofilm samples were used per bacterial strain.

Amplitude sweep and frequency sweep assays were performed in tandem on a single sample, with a total of three samples per bacterial strain. Amplitude sweeps deform the colony biofilms over a range of shear strains while maintaining a constant oscillatory frequency. These measurements can determine the linear viscoelastic region, defined as the range of shear strains for which the measured moduli remain unchanged (**Fig. S1**). Frequency sweep experiments were then performed within the linear viscoelastic region. This involved deforming colony biofilms over a range of rotational oscillatory frequencies, while the amplitude of deformation remained constant. Deformations are proportional to applied forces in the linear viscoelastic region; in this regime, the elastic modulus (G`) and viscous modulus (G``) completely describe the mechanical response of the biofilms.

To prevent moisture loss during the combined amplitude and frequency sweep assays (a total measuring time of around 3 h), a thin layer of paraffin oil was pipetted around the edges of the sample after the plate was lowered to the final position. For amplitude sweep assays, 25 data points were recorded while the oscillating shear strain increased logarithmically from 0.5% - 100% of 1 Hz. The amplitude sweep assays determined that the viscous and elastic moduli did not vary greatly in the 1%-10% shear strain range for all the bacteria we tested (**Fig. S1**), so a shear strain of 5% was used for all frequency sweep measurements. A linear viscoelastic region was not found for the Δ*epsB* mutant, but a shear strain of 5% was used for consistency. For each frequency sweep measurement, 25 data points were recorded while the angular frequency decreased logarithmically from 500 $s^{-1}$ - 0.005 $s^{-1}$.

Cell density and moisture content data was also collected for each sample measured by the rheometer. Cell density was quantified by dilution plating. Samples were serial diluted, plated onto CPG+TZC plates, incubated at room temperature for three days, and colonies were counted. To determine the moisture content of a sample, the wet weight and dry weight of the colony biofilm samples were measured. Wet weight was measured after rheological measurements, and dry weight was measured after samples dried in an oven for at least 4 days at 47-60 °C.

**Bacterial Dispersal and Growth in Tomato Stems**
To quantify bacterial dispersal inside the plant host, we inoculated 36-day-old tomato plants (cv. Moneymaker) with the wildtype *R. pseudosolanacearum* GMI1000 and Δ*epsB* mutant strains. Plants were inoculated following a cut-petiole approach with 2,000 CFU of each bacterial strain (13). At three days post-inoculation, we sampled 100 mg (±10 mg) of tomato stem harvested above and below the inoculation site: +4 cm, +2 cm, 0 cm, -2 cm, -4 cm. Stem samples were homogenized by bead beading. Tissue was added to tubes with 900 µl water and four 2.2 mm metal beads and homogenized using a Qiagen PowerLyzer 24 for two 1.5 min rounds at 2,200 rpm with a 4 min rest. After homogenization, samples were dilution plated, and colonies were counted after two to three days. The detection limit was used for samples that had population sizes below this threshold.

**Microfluidic Biofilm Assay**

Overnight cultures of strains in CPG broth were collected by centrifugation and adjusted to $10^9$ CFU/mL with sterile water. Channels with Polydimethylsiloxane (PDMS) microfluidic devices were pre-coated with 10 mg/mL Carboxymethyl Cellulose-Dopamine (CMC) overnight (7, 14, 15). The adjusted bacterial suspension ($10^9$ CFU/mL) was pumped through the channels until liquid reached the outlet tubing (N= 3 biological replicates). The device was incubated for 6 hours statically (0 µL/hr flow rate) before the channels were flushed and filled with CPG medium. The flow experiment was conducted at 0, 10, 40, 80, and 100 µL/hr flow rates for 72 h. After 72 h, the microfluidic devices were flushed with water to remove planktonic bacteria. Biofilms were stained with Crystal Violet and washed with water. Images were taken at the entry, midpoint, and exit of 10 channels using a compound microscope equipped with an AmScope MD500 camera. Biofilm coverage was measured as percentage of Crystal Violet-stained area over the total channel area examined using ImageJ.

**Biofilm Mobility Assay**

To assess mobility of biofilms, strains were grown on agar, flat (0° angle) or tilted (39° angle). To prepare for the assay, each strain was streaked out from glycerol stocks onto CPG+TZC agar and grown for two days (28 °C) or three days (room temperature) for isolated colonies. Individual colonies were inoculated into CPG broth and incubated overnight at 28 °C and shaken at 250 rpm. Each overnight culture was considered a biological replicate. Each biological replicate was used to prepare two plates, on which three technical replicate droplets of 5 µL of $5x10^4$ CFU/mL cell suspension were pipetted onto CPG+TZC 2% agar in a row near the plate edge. The droplets were allowed to dry in a laminar flow hood. After drying for about an hour, the two plates were placed in a 28 °C incubator on either a flat or tilted surface (0° vs. 39° angle). Plates were incubated for two days at 28 °C before imaged with reflective white light on an Alpha Innotech AlphaImager™ 2200. To enable conversion of pixels to lengths, a ruler was imaged with the camera configuration. Biofilm area was quantified using CellProfiler. Briefly, image illumination was adjusted to make illumination more consistent across the images. A mask was aligned and applied so that the only portion of the image being analyzed was the petri dish. Colony biofilms were identified and the areas were measured. The full CellProfiler analysis pipeline is in Zenodo repository (doi.org/10.5281/zenodo.14767251). Biofilm area was measured for each technical replicate and averaged for the three technical replicates within a plate. For non-RSSC *Ralstonia* strains, images were collected on a BioRad ChemiDoc with either epi- or trans- white light illumination. Strains were categorized as mucoidal based on if their colonies had a glossy appearance or non-mucoidal for dry appearances. Isolates were categorized as having a mobile biofilm based on if their biofilm spread along the agar medium when cultured at a 39° angle.

To further qualitatively assess the mobility individual colonies, agar plates were prepared with inoculum so that approximately 100 colonies would grow per plate. Colonies were grown on CPG+TZC 2% agar plates for three days, either flat (0° angle) or tilted (39° angle). Images were again taken on an Alpha Innotech AlphaImager™ 2200. Like above, a ruler was imaged to convert pixel units to millimeters.

**Area and Population Sizes of RSSC Colonies**

To measure the area of RSSC colonies, agar plates with roughly 500 colonies per plate were cultured and imaged after two days of incubation. Strains were streaked on CPG+TZC agar plates and incubated for two days at 28 °C. Individual colonies were picked and inoculated into CPG broth for each biological replicate (3-4 biological replicates per strain). After overnight incubation at 28 °C, cell density was adjusted to approximately $5x10^3$ CFU/mL, 100 µL was spread on CPG+TZC 2% agar plates, and plates were incubated for 2 days at 28 °C. Plates were imaged in an Alpha Innotech AlphaImager™ 2200 with the reflective white light. To enable conversion of pixels to lengths, a ruler was imaged with the camera configuration.

To quantify colony area, images were processed using CellProfiler (16). Briefly, the illumination of the images was normalized, and masking was applied to constrain the image to the plate. Colonies were identified with an object identification step. Distances between objects were measured and touching objects were excluded from analysis by a filter. The colony area was measured in $pixels^2$ and was converted to $mm^2$ values based on the photograph of the ruler. The CellProfiler analysis pipeline is available in the Zenodo repository (doi.org/10.5281/zenodo.14767251). Several iterations of the pipeline were tested to optimize the distinguishment between single and merged colonies. The pipeline has limitations and did not always identify merged colonies, particularly when three colonies merged into a triangular shape.

A subset of colonies was randomly selected for quantification of the population size per colony. Nine well-isolated colonies per strain were selected from a total of three plates. The colony and the underlying agar were excised with a sterile scalpel with care to avoid disturbing the colony while minimizing the volume of agar. Colonies and agar sections were homogenized in separate tubes with 1 mL water and 3 sterile 2.38 mm metal beads. Tubes were shaken at 2,200 rpm for

2 min in a Qiagen PowerLyzer 24. The homogenate was dilution plated, and colonies were counted after three days of incubation at room temperature.

**Growth Curve Assay**

Growth curves were measured for strains in CPG broth, iBMM with 10 mM glucose, or iBMM with 10 mM glutamate to determine differences of fitness in liquid media. Strains were streaked for isolation onto CPG+TZC agar from glycerol stocks. For each biological replicate, individual colonies were inoculated in CPG broth and incubated overnight. Overnight cultures were washed three times using 1 mL of water in which cells were pelleted at 10,000 rcf for 1 min, the supernatant was removed, and the pellet was resuspended. After washing, densities were normalized to $2.5 \times 10^7$ CFU/mL using $A_{600}$ measurements. To prepare a 96-well plate for the growth curve measurements, 20 µL of the density normalized suspensions were added to 180 µL media and plates were sealed with a Breath-Easy sealing membrane. Growth curves were measured in a BioTek Synergy H1 Plate Reader with $A_{600}$ measurements every 1 hr, continuous, orbital shaking at 28 °C. Growth curve data was collected using Gen5 software and analyzed using Growthcurver (17).

**Evolutionary Analyses of RSSC Biofilm Factors**

The presence/absence of 17 EPS-I genes, *xpsR*, *nucA,* and *nucB* were determined for 399 RSSC genomes and 72 genomes of other species in the *Ralstonia* genus. We performed BLASTp searches (18) using KBase (19), typically using 40% sequence identity and 50% alignment overlap thresholds. The *epsC* gene encodes a WecB-family enzyme that has a second paralog in some RSSC genomes, which may be the poorly characterized "*ops*" cluster that is implicated in O-antigen biosynthesis and EPS-I production in *Rsol* K60 (20). To distinguish *epsC* from the putative *ops* paralog, a multiple sequence alignment (MSA) was created using MUSCLE (21) with default KBase parameters. An approximate maximum likelihood gene tree was created using FastTree2 (22) with default KBase parameters, and the tree was used to distinguish *epsC* from the paralog. Results from the BLASTp analysis were visualized on a KBase SpeciesTree (v2.2.0, default KBase parameters) in iTOL (23).

For the gene neighborhood analysis, we downloaded genomic regions surrounding the genes-of-interest in GenBank flat format (.gbk) from the NCBI Graphics Viewer. We ran Clinker (24) using these .gbk files, which yielded interactive synteny files with links between genes based on global amino acid identity.

To further study the evolutionary history of the RSSC *eps* cluster, we conducted synteny and phylogenetic analysis of *eps* genes referencing a database of 16,355 Betaproteobacteria genomes obtained using *Mycotools* v0.30.36 (25). We examined the shared synteny and gene conservation of the EPS cluster across this dataset using the Cluster Reconstruction and Phylogenetic Analysis Pipeline (CRAP) implemented in *Mycotools* (26). Briefly, CRAP identifies homologs of *eps* genes using *blastp* (27)*;* truncates large homolog sets using iterative sequence similarity clustering via *mmseqs cluster* to yield a maximum of 1000 homologs (28); reconstructs phylogenies using *mafft* alignment (29), *clipKIT* trimming (30), and *fasttree* reconstruction (22); imposes synteny diagrams of extracted loci onto the tips; and extracts the most similar locus to the query for each genome via Jaccard similarity of gene overlap. Gene presence-absence for each genome is based on the outputted loci generated from CRAP. To determine if the evolution of acetyltransferase genes is congruent with species evolution, we reconstructed phylogenomic trees for genomes containing RS_RS21980 homologs. These phylogenomic trees were generated by identifying single-copy orthologs using *db2hgs* implemented in *Mycotools* and reconstructing a multigene partition phylogenomic tree using *IQ-TREE2* with 1,000 ultrafast bootstrap replicates.

As a third approach, the tool cblaster was used to search the *eps* gene cluster against NCBI nr. cblaster performs BLAST based searches to identify similar gene clusters. We only included the 17 genes that encode biosynthesis or export functions and did not include the *xpsR* regulator in the cluster search. cblaster performs individual BLAST searches against NCBI nr for each sequence in the query. Then cblaster uses the NCBI Identical Proteins database to identify the genetic coordinates of each hit and will return gene clusters that have the user-specified number of homologous genes within the user-specified genetic distance. We required the search to identify 10 homologs of the 17 *eps* genes within 20 kb of the neighboring genes. cblaster identified matching clusters in only RSSC genomes (n=507) including the *R. syzygii* genomes labeled as the pre-2015 "Blood Disease Bacterium" name. The full settings for the search and the results are stored in the Zenodo repository (doi.org/10.5281/zenodo.14767251).

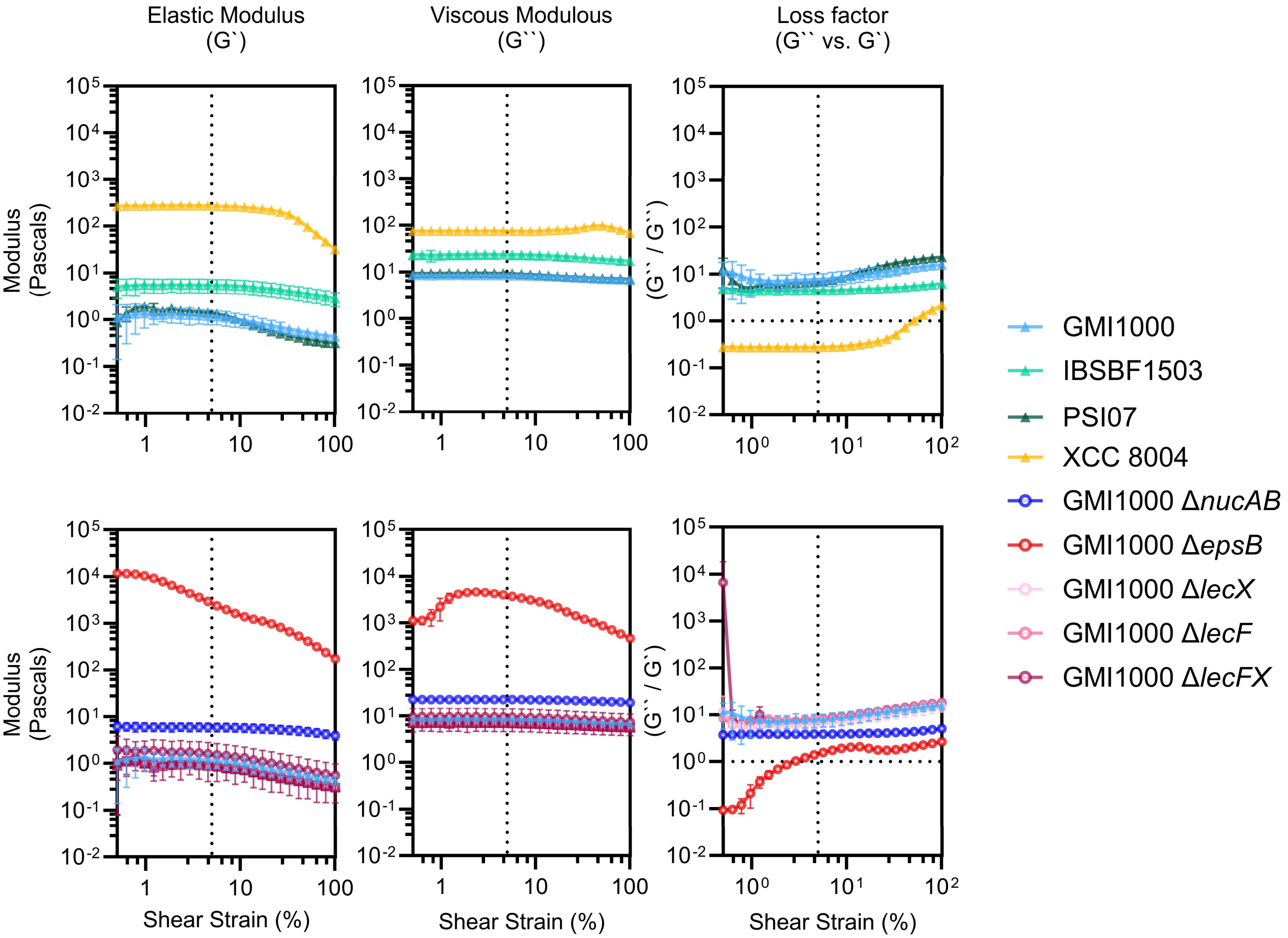


**Fig. S1. Linear viscoelastic regions of samples were determined with amplitude sweep measurements.** Viscous and elastic moduli were measured over increasing shear strain, and the linear viscoelastic regions were determined for all samples except for the GMI1000 Δ*epsB*. The vertical dotted line at a shear strain of 5% indicates the chosen shear strain for all subsequent frequency sweep experiments. In the loss factor plots, the horizontal dotted line indicates the value when the viscous and elastic moduli are equivalent. The symbols represent the mean, and the error bars represent the standard deviation. Three biological replicates for each bacterial strain.

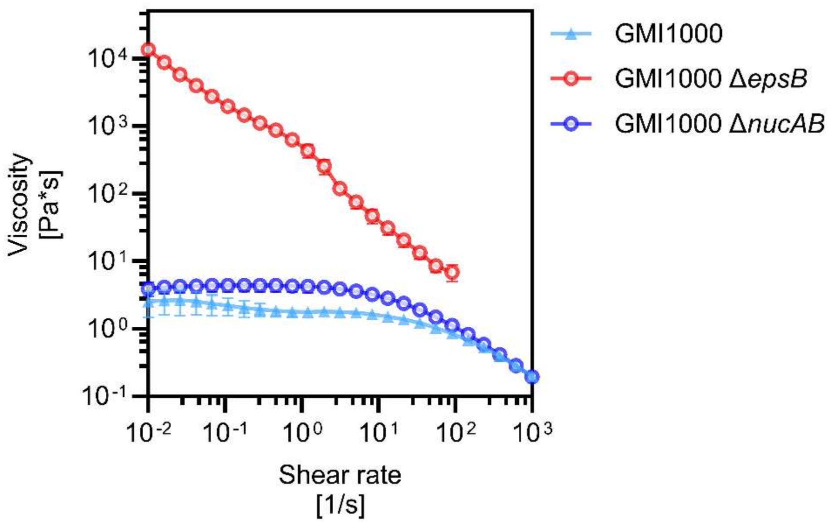


**Fig. S2. The viscosity of GMI1000 wild type, GMI1000 Δ*epsB*, and GMI1000 Δ*nucAB* measured at increasing shear rates from 0.01 $s^{-1}$ to 1000 $s^{-1}$.** The GMI1000 Δ*epsB* samples were only measured at shear rates ≤ 100 $s^{-1}$ because at higher shear rates, the samples formed air gaps between the rheometer measuring tool and base plate. The wildtype GMI1000 data is repeated from **Fig. 1D** for comparison to the mutants. Data for the Δ*lec* mutants can be found in Carter *et al*. 2024 (11). The symbols represent the mean, and the error bars represent the standard deviation. Three biological replicates for each bacterial strain.

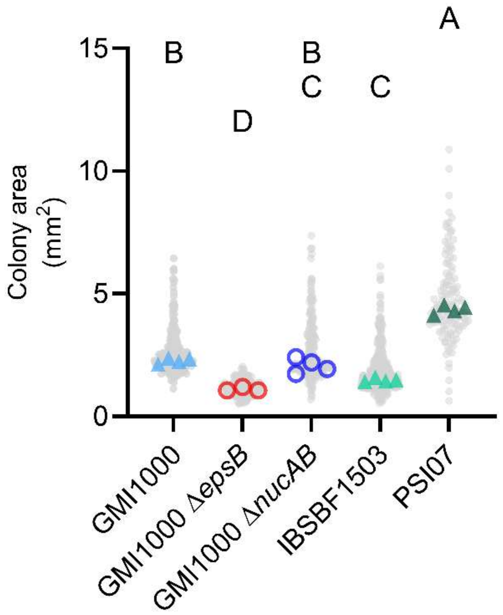


**Fig. S3**. **Sizes of individual colonies of RSSC wildtype and mutant strains grown on rich agar.** Data from GMI1000 wildtype and Δ*epsB* mutant are repeated from **Fig. 3D**. Colored points represent the median colony size per replicate agar plate, and grey points represent the colony size of individual colonies measured. Statistical groupings are based on a Welch ANOVA test and a Brown-Forsythe multiple comparisons test of the medians of each replicate ($p < 0.05$).

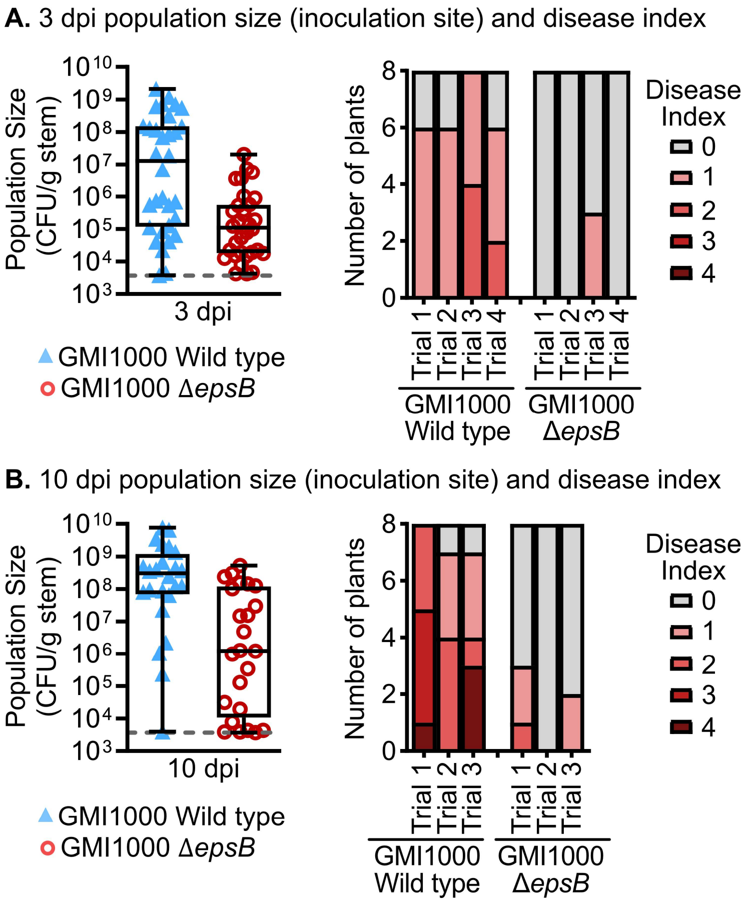


**Fig. S4. The Δ*epsB* mutant has a growth defect and a virulence defect in tomato plants.** Tomato cv. Moneymaker plants were cut-petiole inoculated with GMI1000 wild type or the Δ*epsB* mutant. Left panels show population sizes at the site of inoculation at (***A***) 3 or (***B***) 10 days post inoculation (dpi) for 3-4 trials of n=8 plants per strain; the 3 dpi data is repeated from **Fig. 3A** for comparison. Right panels show the disease index of the plants when they were destructively sampled: 0 corresponds to 0% leaves wilted, 1 corresponds to ≤25%, 2 corresponds to ≤50%, 3 corresponds to ≤75%, and 4 corresponds to ≤100%.

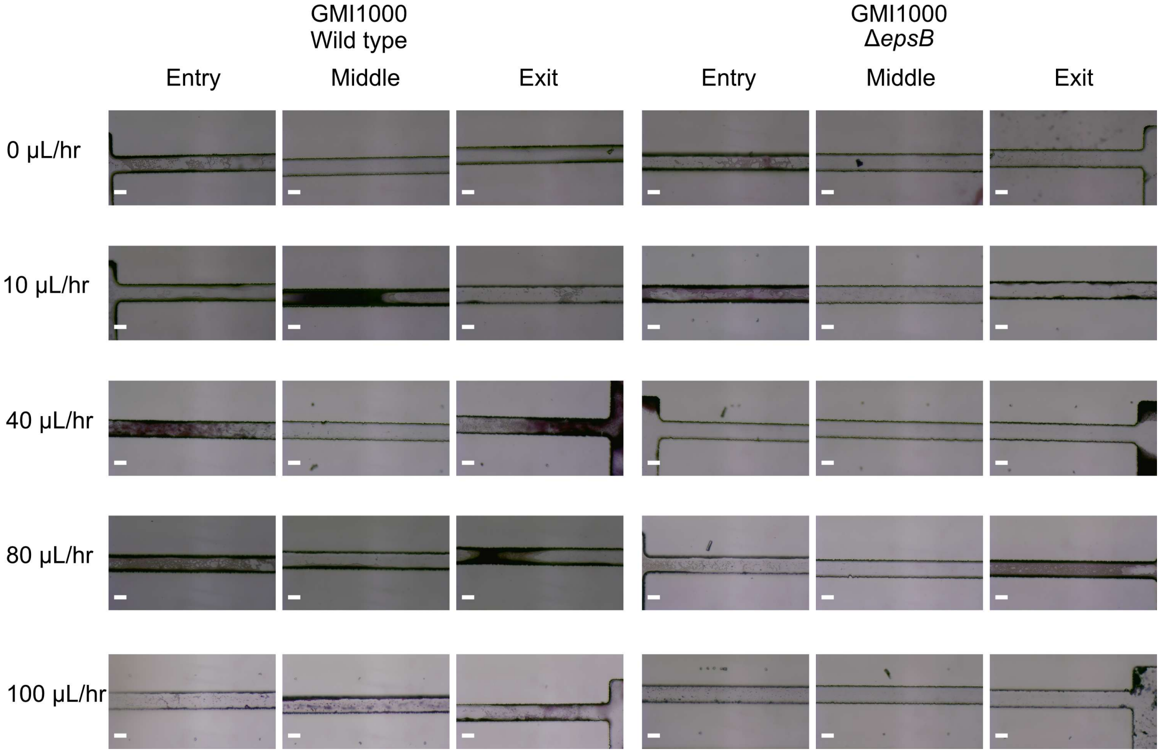


**Fig. S5. The Δ*epsB* was less efficient in colonizing microfluidic channels under flow conditions.** CMC-Dopamine-coated microfluidic channels colonized with wildtype GMI1000 and GMI1000 Δ*epsB* and subjected to various flow rates of rich broth (CPG) for 72 hours. Channels were washed with water to remove planktonic cells and stained with Crystal Violet prior to imaging (Bar, 0.1 mm). The images are representative of three biological replicates.

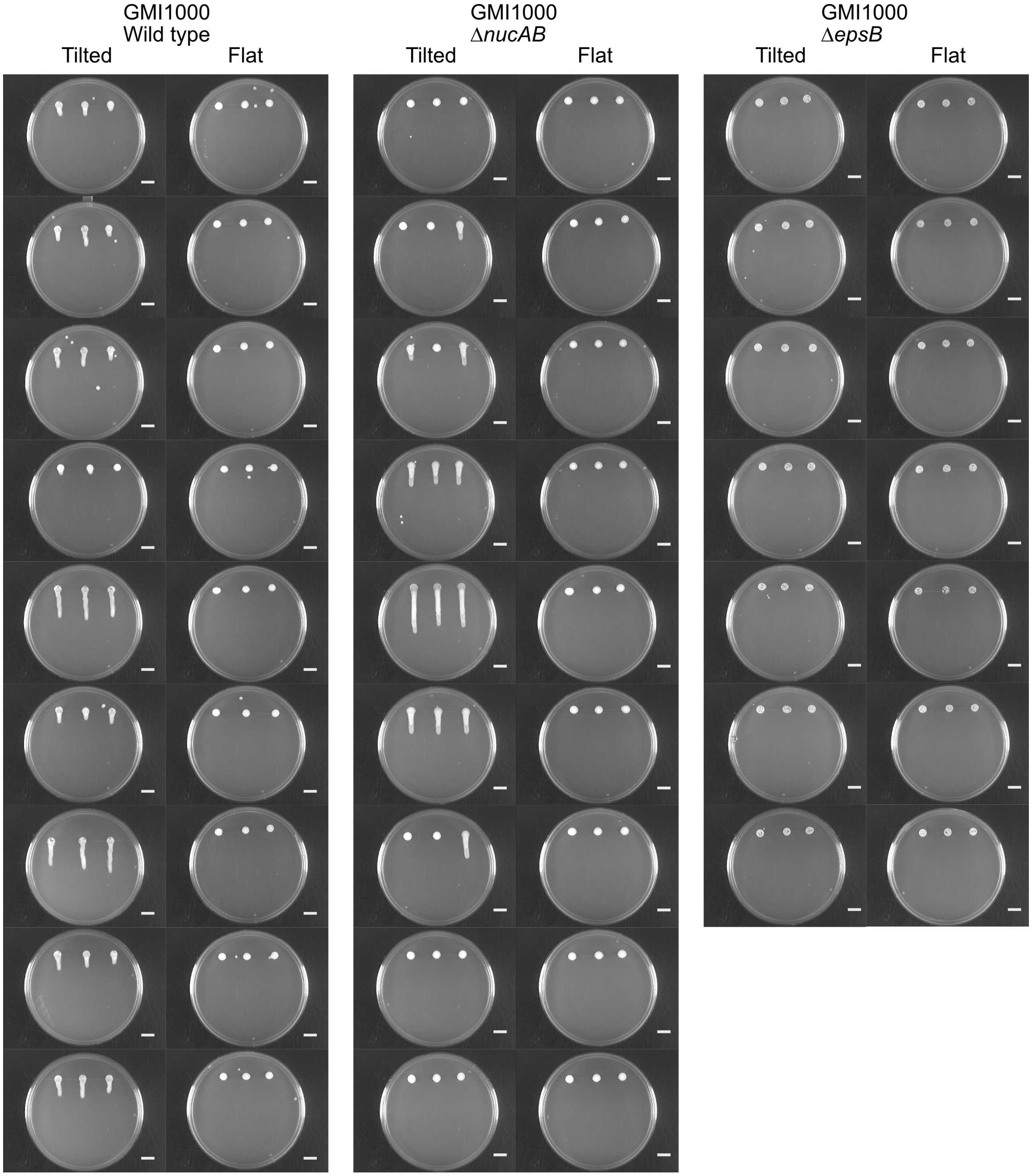


**Fig. S6. Images of RSSC biomass when grown at a flat or tilted at a 39˚ angle.** Strains were inoculated on rich agar plates at a normalized density of 125 CFU/spot and grown at their respective angles for two days. Flat and tilt images are paired so that side-by-side images show colony biofilms grown from the same liquid culture. Spots on the same plate were treated as technical replicates (scale bar, 1 cm).

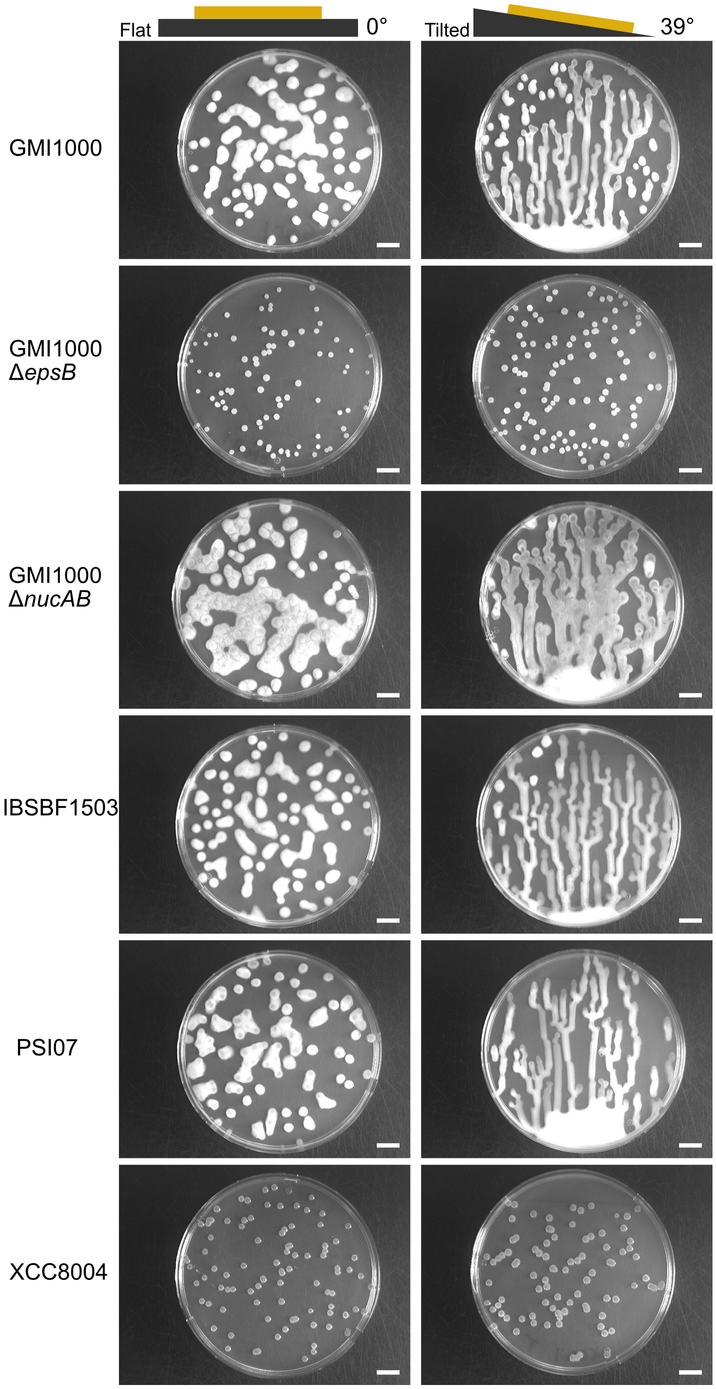


**Fig. S7. Colony mobility behavior of Wildtype RSSC strains, RSSC mutants (GMI1000 Δ*nucAB* and GMI1000 Δ*epsB*), and wildtype *Xanthomonas* XCC8004.** Strains were inoculated to yield single, isolated colonies by spread-plating a diluted bacterial culture with glass beads on the agar medium. GMI1000 Δ*epsB* and XCC8004 colonies were mobile. Colonies grew for three days at a constant flat angle or a constant tilt of 39˚. The plates were paired, so side-by-side images represent colonies grown from the same liquid culture. Representative photos shown for six biological replicates. The scale bar corresponds to 1 cm.

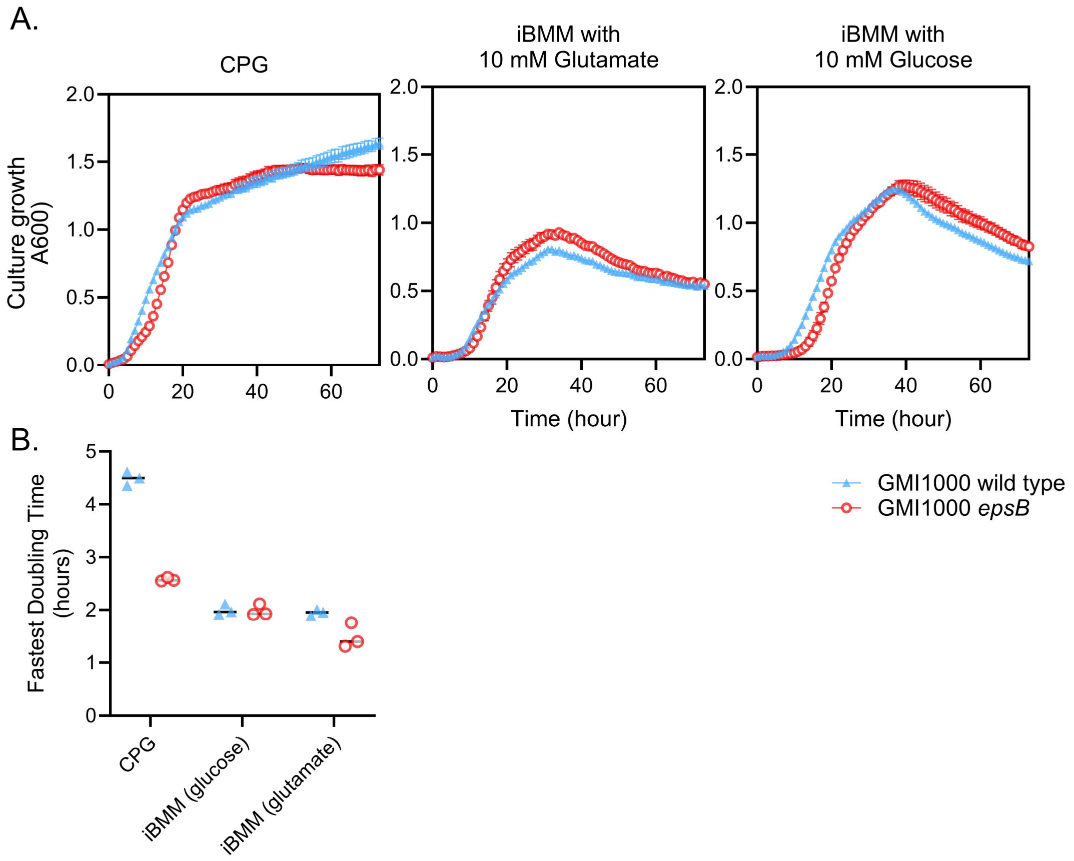


**Fig. S8. Growth of GMI1000 wild type and Δ*epsB* mutant in liquid media.** ***(A)*** Cultures were grown at 28 °C with constant shaking in a BioTek plate reader in rich CPG medium or improved Boucher's minimal medium (iBMM) pH 6.5 with 10 mM glucose or glutamate. The symbols represent the mean, and the error bars represent the standard deviation. Three biological replicates for each bacterial strain. ***(B)*** Fastest doubling time calculations were determined using Growthcurver (30). The greatest difference in the growth rate between GMI1000 Δ*epsB* and the wild type was observed in CPG broth, where the Δ*epsB* mutant can growly nearly twice as fast as the wild type at their max growth rates. Bars represent the median.

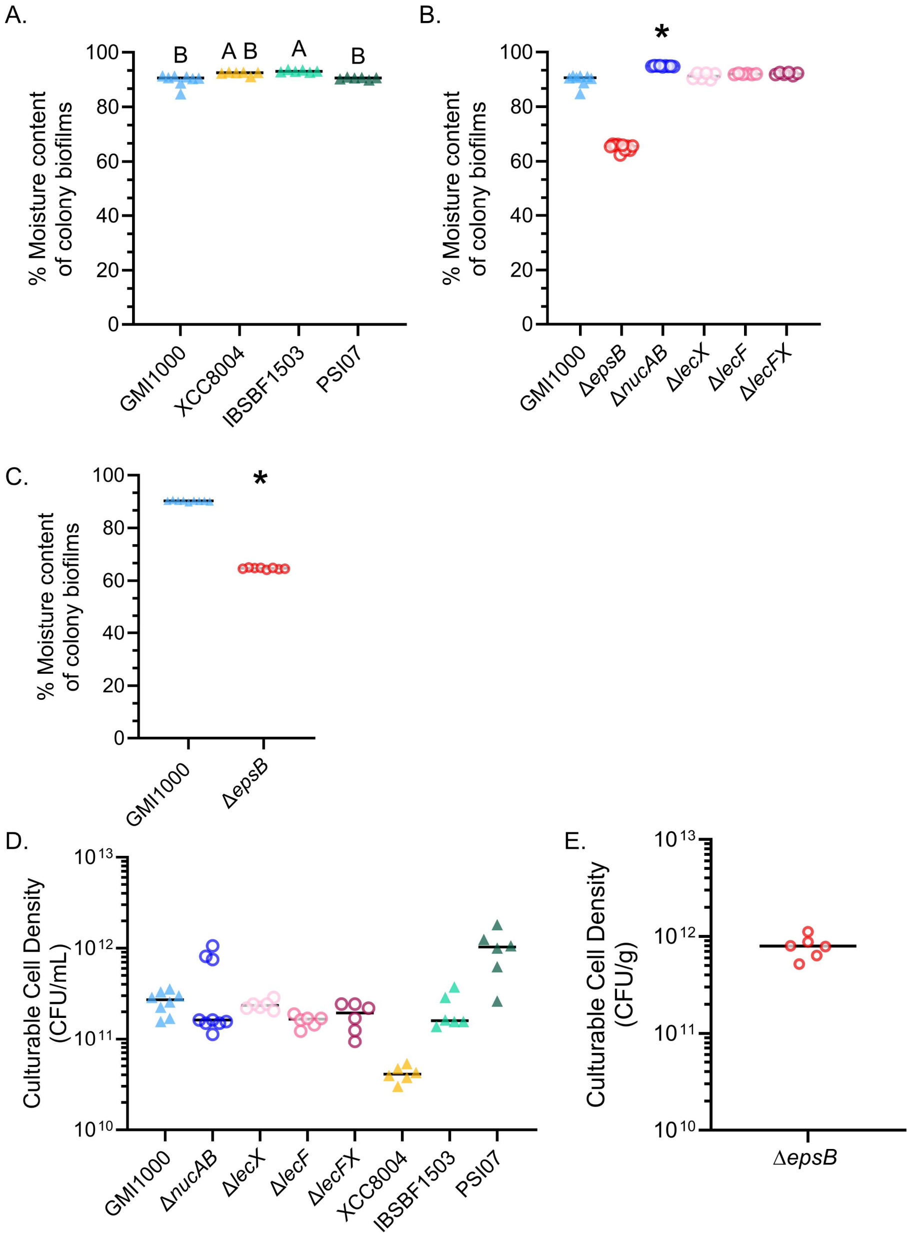


**Fig. S9. Cell density and moisture content of colony biofilm samples varied, and GMI1000 Δ*epsB* colony biofilms were drier than the wild type.** ***(A-B)*** Moisture content of wildtype and mutant colony biofilms. Wildtype GMI1000 data is repeated across in panel A and panel B for clear comparisons. Moisture content was measured in the exact colony biofilm samples used for each rheometry experiment. Thus, each sample was not measured on the same day. Bars represent the median. ***(A)*** *Rsol* IBSBF1503 moisture content was significantly higher than the other two RSSC wild types (Letters indicate $p < 0.05$ by Kruskal-Wallis Multiple Comparisons Test; All vs. all). ***(B)*** The Δ*nucAB* mutant had increased moisture content (* indicates $p < 0.05$ by Kruskal-Wallis and Dunn's Multiple Comparisons Tests; GMI1000 vs. all). ***(C)*** Moisture content measurements were repeated to directly compare the moisture content of wildtype GMI1000 and the Δ*epsB* mutant. For the initial rheology experiments, an early sample of wildtype GMI1000 was grown on older agar plates. The sample from aged plates had lower biofilm moisture than all subsequent biofilms grown on freshly poured agar plates at 84.7% and 88.7% moisture content. We chose to not remove this sample as an outlier. However, this decision meant that the GMI1000 biofilm moisture data was not-normally distributed, and comparisons of GMI1000 vs. Δ*epsB* mutant with the non-parametric Kruskal-Wallis and Dunn's multiple comparisons test yielded p = 0.371 despite the trend. Thus, we repeated the experiment using the same batch of freshly poured plates to grow colony biofilms for moisture content measurements. When the moisture content of wildtype GMI1000 and Δ*epsB* colony biofilms grown on the same batches of media were compared, the difference was consistent and significantly different, and the variation of wildtype GMI1000 samples decreased (* indicates $p < 0.0001$ by unpaired T test). We report both datasets for transparency and to also reinforce the importance of agar plate conditions for colony moisture and rheology conditions. Bars represent the mean.

***(D-E)*** Cell densities of colony biofilm samples used in rheometry experiments. Bars represent the median. ***(D)*** Cell densities of most genotypes were measured by pipetting colony biofilm samples into 1 mL of water, homogenizing, and dilution plating to get CFU/mL measurements. ***(E)*** Because GMI1000 Δ*epsB* samples were too viscous to be pipetted, a portion of the sample was transferred into a tube, weighed, homogenized in water, and dilution plated to get CFU/g measurements.

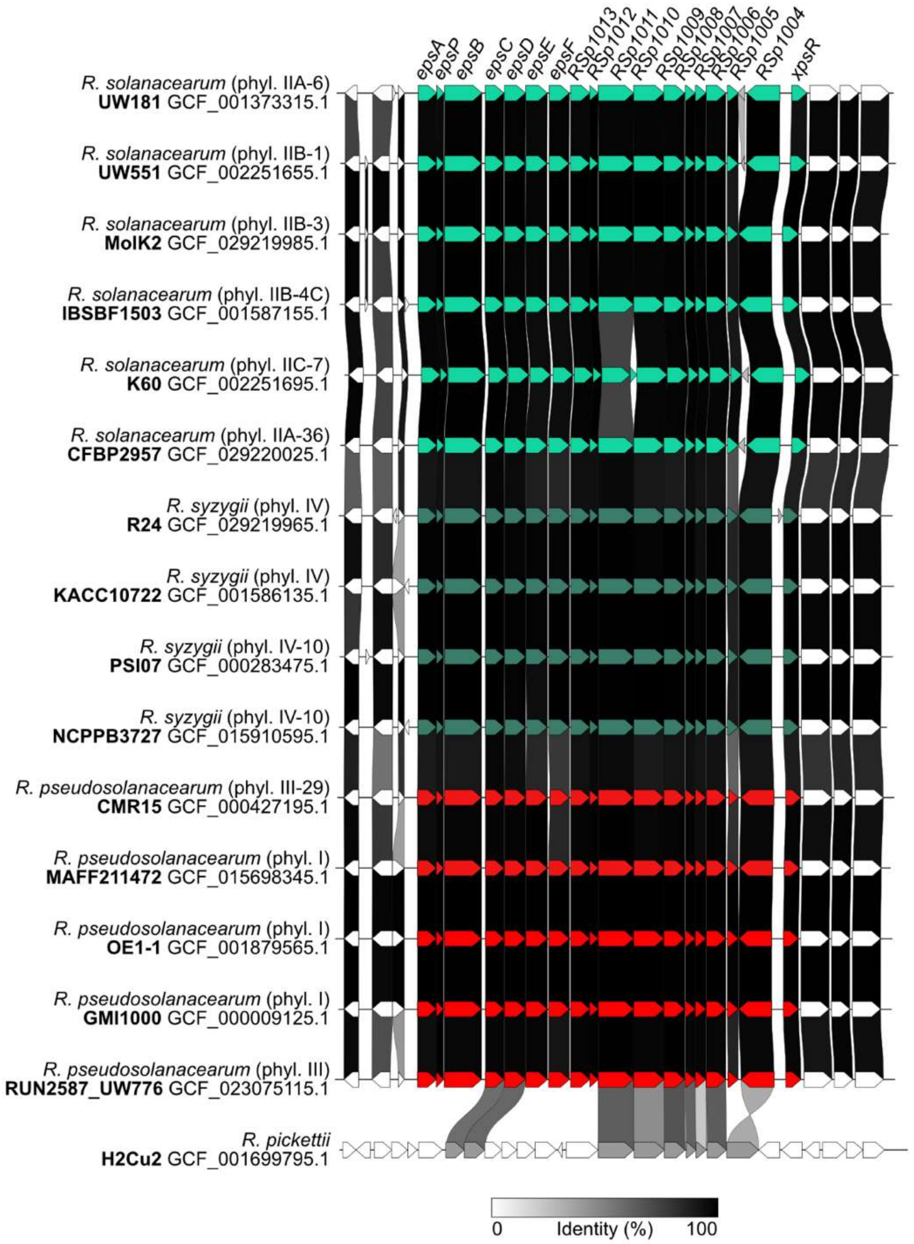


**Fig. S10. Synteny of the *eps* gene cluster is broadly conserved across phylogenetically diverse RSSC.** The 17 EPS-I production genes and the *xpsR* regulator are labeled with the gene name or GMI1000 locus tags. For each strain, the strain name is bold and the label includes the NCBI genome accession, species name, and phylotype-sequevar. In the initial BLASTp analysis, one non-RSSC genome, *Ralstonia pickettii* H2Cu2 encoded BLAST hits to 9 of the 17 EPS-I production genes. The relatively low identity and the absence of many *eps* homologs suggests that this genome encodes a distinct cluster with divergent biosynthesis. Visualizations were created in Clinker, which draws linkages between homologous genes that represent the pairwise global amino acid identity percentage.

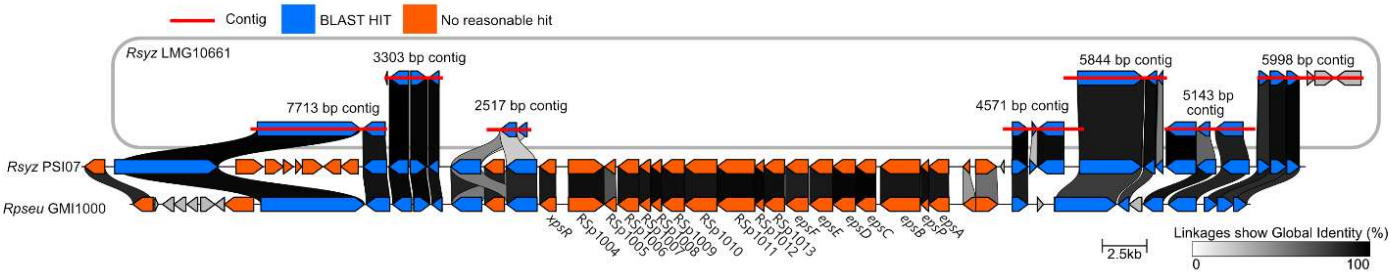


**Fig. S11. Investigation of the RSSC genome (GCF_919592095.1) that lacks BLAST hits for the *eps* genes.** The genome is assembled into 348 contigs to a total size of 4.0 Mb. Because most RSSC genomes exceed 5 Mb, the genome is suppressed on RefSeq with the warning “genome length too small”. BLAST and Clinker were used to investigate the presence of the *eps* gene neighborhood in GCF_919592095.1. Blue and orange show genes that respectively matched or lacked homology to contigs in GCF_919592095.1; red lines indicate the full length of the matching contigs. Interpretation: Although the absence of EPS BLASTp hits in this fragmented genome could be due to assembly errors, this genome belongs to a strain in an insect-transmitted, genome-reduced lineage of *R. syzygii* that causes Sumatra disease of clove in Indonesia. The original taxonomic definition of the Sumatra disease of clove pathogens describes these strains as varying in their colony morphology from “viscid” to “butyrous” (31). Butyrous is the common adjective for the dry colonies that are produced by RSSC mutants that do not produce EPS-I. In imperfect culture storage, RSSC strains can readily “phenotype convert” from mucoid to dry/butyrous colonies, and the strains analyzed in the taxonomy study had been stored in Indonesia and shipped to the United Kingdom before analysis. The original isolation report of the Sumatra disease pathogen describes unique aspects of these pathogens’ fastidious growth and colony morphology, but they do not report butyrous variants (32). Thus, it remains unclear as to whether there are any rare RSSC wild types that naturally lack the *eps* genes.

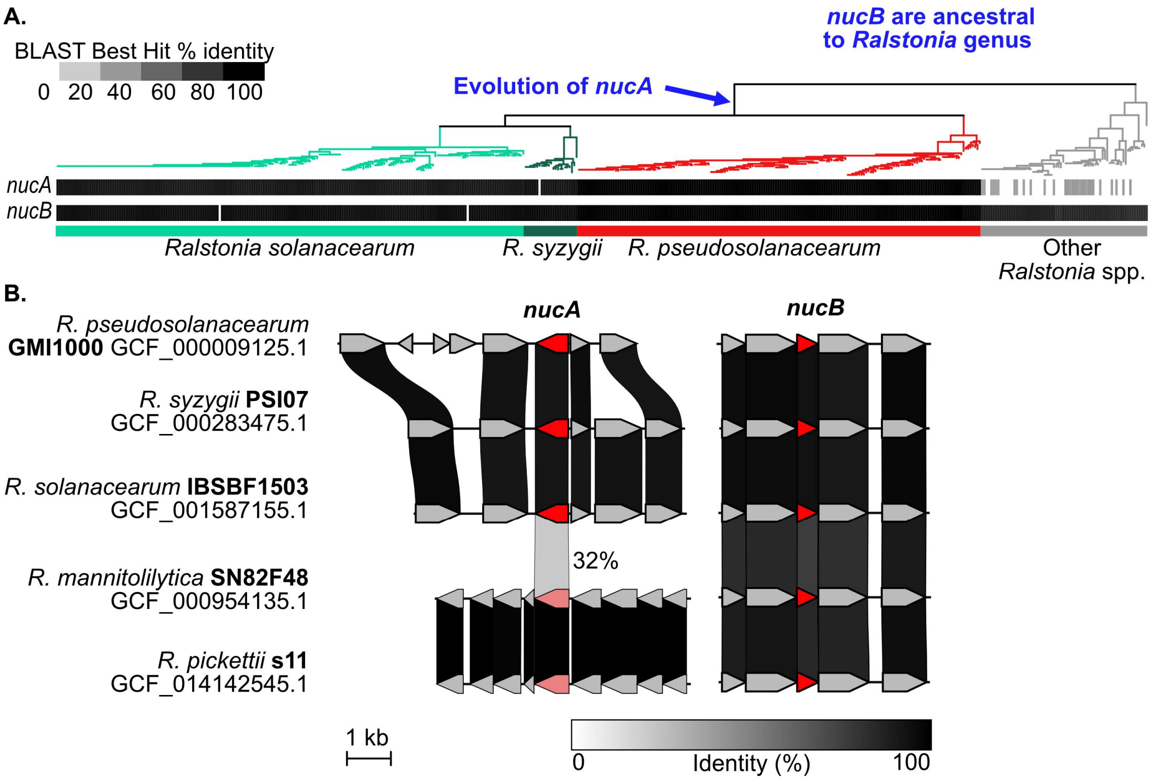


**Figure S12. Inferred evolutionary history of the extracellular nucleases, NucA and NucB.** ***(A)*** Presence of *nucA* and *nucB* genes in genomes of three species of *Ralstonia* wilt pathogens and in genomes of ten *Ralstonia* species that do not wilt plants was determined by BLASTp queries on the KBase Platform. Rectangles show percent amino acid identity of the best BLAST hit for queries with *R. solanacearum* K60 proteins. The approximate maximum likelihood phylogenetic tree was built from 49 conserved bacterial genes using the KBase Insert Genomes into SpeciesTree v2.2.0 application and visualized in iTOL. High identity NucB homologs were identified in most *Ralstonia* genomes whereas high identity NucA homologs were only identified in RSSC genomes. ***(B)*** Synteny analyses of the gene neighborhoods around *nucA* and *nucB* homologs. The high identity NucB homologs were encoded in the same genomic context across the *Ralstonia* genus. RSSC NucA homologs were encoded in a consistent genomic location while low-identity BLASTp hits from non-RSSC genomes were encoded in a divergent genomic location. Blue text indicated the hypothesized evolutionary history of the nucleases where *nucB* was ancestral to the entire genus and *nucA* was acquired by the ancestor of the RSSC lineage.

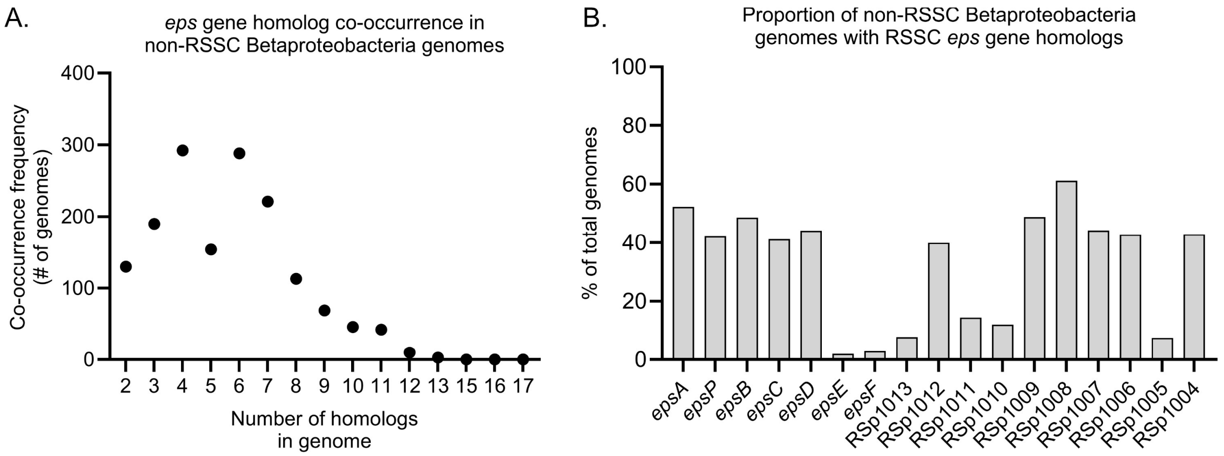


**Fig. S13. *eps* gene homologs are rare amongst the Betaproteobacteria.** The cluster reconstruction and phylogenetic analysis pipeline was used to search genomes of 16,355 betaproteobacteria for two-or-more homologs of RSSC *eps* genes within 20 kb, yielding 1,563 hits of non-RSSC genomes. (A) The number of *eps* homologs in each genome. (B) Relative abundance of each *eps* gene family in non-RSSC betaproteobacterial genomes that have at least 2 clustered *eps*-like genes. The least common *eps* genes were *epsE*, *epsF*, RSp1013, RSp1011, RSp1010, and RSp1005.

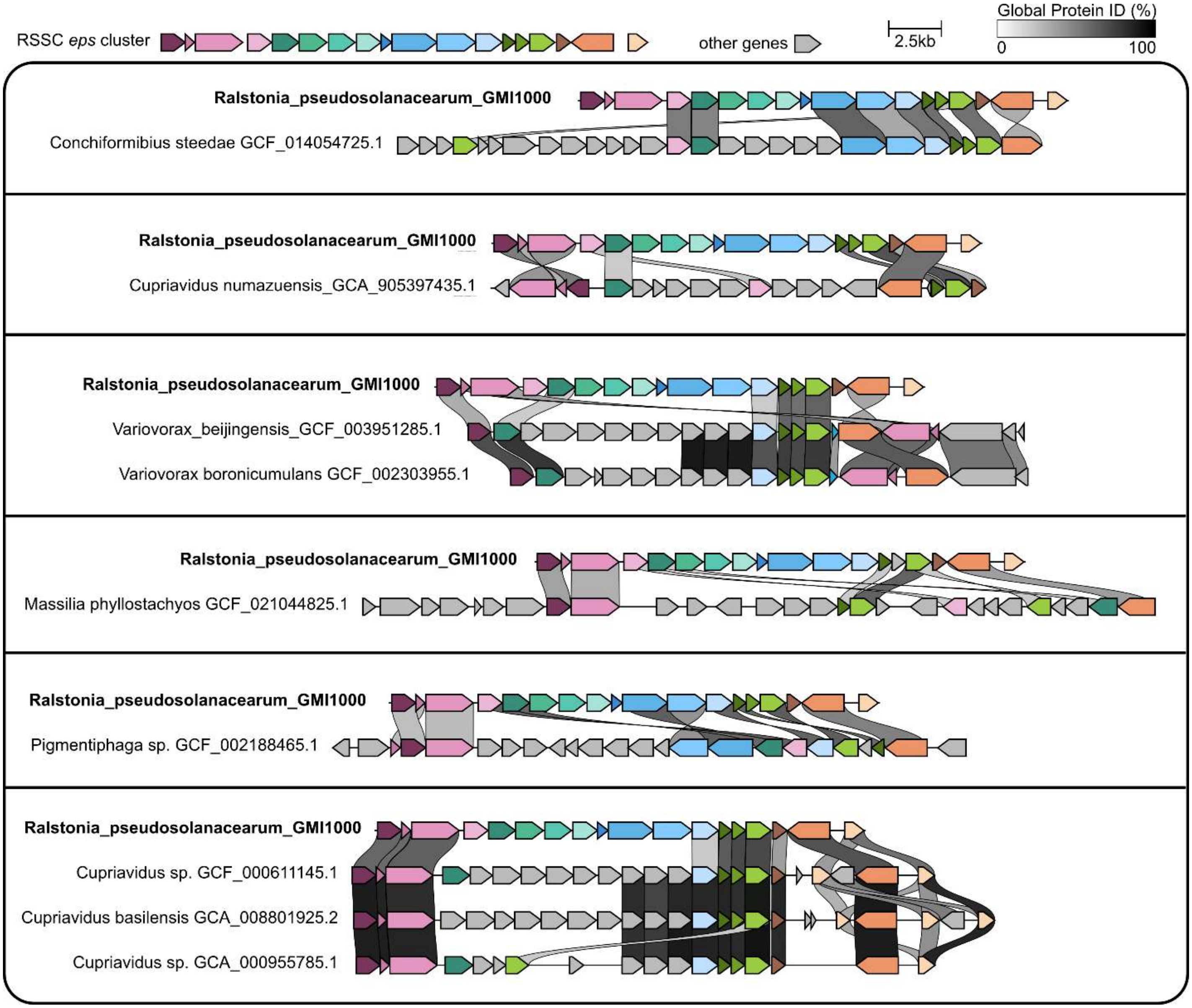


**Fig. S14. Genetic organization of exopolysaccharide gene clusters identified in diverse Betaproteobacteria.** The genomes from the cluster reconstruction and phylogenetic (CRAP) analysis with the most homologs of RSSC *eps* genes were visualized with Clinker.

Homologs to RSSC *eps* genes are shown in distinct shades of pink, teal, blue, green, or orange. Grey indicates genes that were exclusively found in non-RSSC gene clusters.

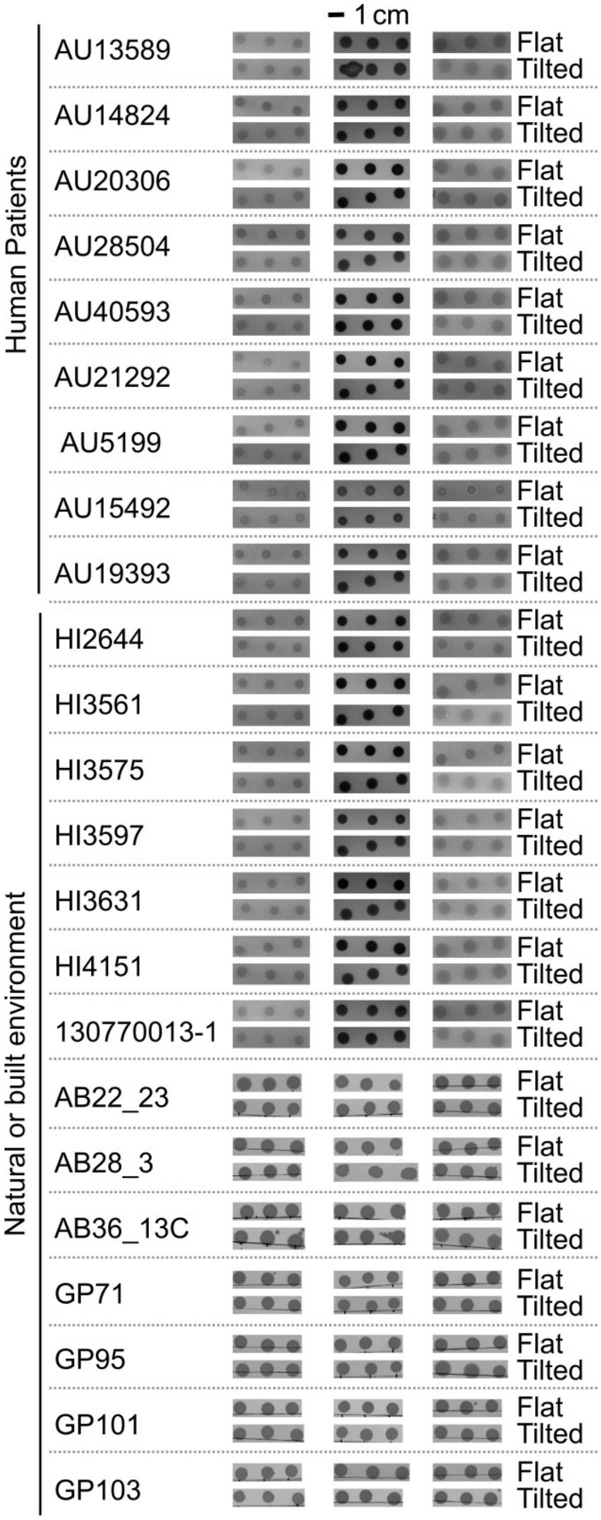


**Fig. S15. Colony biofilms of non-RSSC *Ralstonia* lack mobility**. Strains were spotted at a normalized density of 125 CFU/spot on rich agar and plates were incubated on either a flat surface or a surface tilted at 39°. Three biological replicate spots were included in each of the 3 experimental trials. The scale bar is 1 cm.

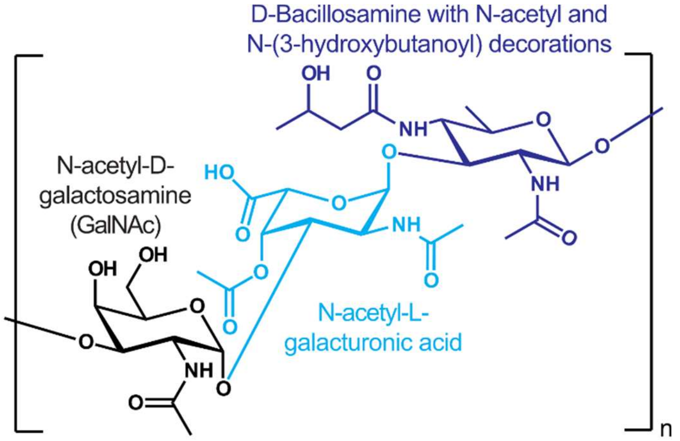


**Fig. S16. The chemical structure of the EPS-I polymer produced by *Ralstonia solanacearum* species complex bacteria.** The structure was determined in Orgambide *et al.* 1991 (33).

**Table S1.** Biofilm rheology literature analysis.

| Species/Sample type | Microbial biofilm (B), Polymer extract (P), or other (O)? | Lifestyle or environment of organism | Culturing method prior to rheology | G` > G``(within the linear viscoelastic region) | Reference | Comments |
|---|---|---|---|---|---|---|
| *Candida albicans* | B | commensal/opportunistic pathogenic yeast (fungus) | colony biofilms on polycarbonate filters on Spider agar; grown 1 week and passaged to a new plate for an additional week | yes | (34) | |
| Mixed culture biofilm of yeasts: *Rhodototula mucilaginosa, Candida krusei, Candida kefyr, Candida tropicalis* | B | Yeast isolated from polyvinylidene–fluoride ultrafiltration (UF) membranes from a clarified apple juice processing facility | (1) static biofilms grown in filter-sterilized apple juice; cells and juice replenished every 72 hours for 3-11 weeks and (2) a turbulent flow bioreactor (Re > 200,000) with cells and juice replenished every 72 hr for 8 weeks | yes | (35) | |
| *Mycobacterium abscessus* (both colony biofilm variants) | B | opportunistic lung pathogen (esp. in cystic fibrosis patients) | colony biofilms on nitrocellulose membranes on 7H10 agar | yes | (36) | Rough and smooth colony biofilm variants tested, both were elastic dominant |
| *Escherichia coli* | B | animal gut microbe | | yes | (37) | Curli proteins have the largest impact on colony viscosity |
| Mixed Culture Biofilm | B | surface of nano-filtration (NF) membranes from a drinking water plant | Natural biofilms scraped from nano-filtration membranes used for years in a water treatment plant | yes | (38) | |
| *Comamonas denitrificans* | B | Denitrifier in Birtley Wastewater Treatment plant | colony biofilms on Tryptic Soy Agar (1.5% agar) | yes | (39) | |
| *Pseudomonas aeruginosa* | B | opportunistic pathogen | colony biofilms on Nutrient Agar (1.5% agar) | yes | (39) | |
| *Bacillus subtilis* | B | Rhizosphere and soil microbe | colony biofilms on LB Miller Agar (1.5% agar) | yes | (39) | |
| *Pseudomonas fluorescens* | B | isolated from prefilter tanks (in water treatment plant?) | colony biofilms on Nutrient Agar (1.5% agar) | yes | (39) | |
| *Bacillus subtilis* | B | Rhizosphere and soil microbe | Biofilms grown on sand. | Yes (mostly, see comment) | (40) | Surfactin reduces and EPS increases viscoelasticity; eps deficient mutant appeared to be viscous dominant, however it was |

| Species/Sample type | Microbial biofilm (B), Polymer extract (P), or other (O)? | Lifestyle or environment of organism | Culturing method prior to rheology | G` > G``(within the linear viscoelastic region) | Reference | Comments |
|---|---|---|---|---|---|---|
| | | | | | | also the noisiest dataset |
| *Serratia marcescens* | B | opportunistic, catheter-associated pathogen | Colony biofilms grown for 7 days on LB agar with varying agar concentration (1%, 1.5%, 1.75%, and 2% w/v) | yes (though it does not seem that the linear viscoelastic region was measured even if stated in methods) | (41) | Linear viscoelastic region approximated at 2 % strain, but strain sweep plot does not appear to be linear around this strain except for the measurements of samples from the 1 % agar condition. |
| *Pseudomonas aeruginosa* (natural variants evolved within cystic fibrosis patients' lungs) | B | | Colony biofilms grown overnight on agar | yes | (42) | Differentiated the impact of Psl, Pel, and alginate on biofilm rheology. More alginate is known to cause "alginate softening of biofilms" (lower elasticity). Increased Psl stiffens biofilm (probably in combination with cross-linking by the CdrA protein), and Pel increases the ductility of the biofilm. |
| *Bacillus subtilis* | B | soil bacteria | biofilm grown on agar media | yes | (43) | |
| *Azotobacter vinelandii* | B | soil bacteria | biofilm grown on agar media | yes | (43) | |
| *Pseudomonas aeruginosa* | B | opportunistic pathogen | colony biofilms grown on LB with 1.5% w/v agar; variations in metals added | yes (mostly, at high frequencies there are two points where the biofilms are viscous dominant) | (44) | FeCl3 and Al2(SO4)3 increased and citrate reduced biofilm viscoelasticity. |
| *Stenotrophomonas maltophilia* | B | Opportunistic pathogen | Colony biofilm on polycarbonate filters on R2A agar | yes | (45) | |

| Species/Sample type | Microbial biofilm (B), Polymer extract (P), or other (O)? | Lifestyle or environment of organism | Culturing method prior to rheology | G` > G``(within the linear viscoelastic region) | Reference | Comments |
|---|---|---|---|---|---|---|
| *Staphylococcus epidermidis* | B | skin microbe that is an opportunistic, catheter-associated pathogen | biofilms grown in bioreactor with an in situ rheometer | yes | (46) | |
| *Pseudomonas aeruginosa* | B | | Pellicle biofilms (formed at liquid-air interface) grown in LB | yes | (47) | |
| *Escherichia coli* | B | commensal, non-pathogenic, gut strain | colony biofilms on salt-free LB agar (1.8% agar) | yes | (48) | |
| Mixed Culture Biofilm | B | natural seawater at Newcastle University's Dove Marine Laboratory | static biofilms formed on glass coupons incubated in natural seawater over two months | yes | (49) | |
| Mixed culture biofilm (*Pinnularia* spp.) | B | Brown microalgae from river | Sampled from rivers in Spain and microbes identified by FISH (fluorescent in situ hybridization) | yes | (50) | |
| Mixed culture biofilm (*Euglena mutabilis*) | B | Photosynthetic protist from river | Sampled from rivers in Spain and microbes identified by FISH (fluorescent in situ hybridization) | yes | (50) | |
| Mixed culture biofilm (*Chlorella* spp.) | B | Green microalgae from river | Sampled from rivers in Spain and microbes identified by FISH (fluorescent in situ hybridization) | yes | (50) | |
| Mixed culture biofilm (*Leptospirillum* spp. and *Acidiphilium* spp.) | B | Mixed bacterial biofilm from river | Sampled from rivers in Spain and microbes identified by FISH (fluorescent in situ hybridization) | yes | (50) | |
| Mixed Culture Biofilm | B | water from Montana State University duck pond | biofilms grown on the spinning disk rheometer plates in a flowing bioreactor | N/A | (51) | Elastic moduli shown but not viscous moduli in table 1 |
| *Streptococcus mutans* | B | dental pathogen | biofilms grown on rheometer disk plates in flowing bioreactor | yes | (52) | |
| *Streptococcus mutans* | B | dental pathogen | biofilms grown on the spinning disk rheometer plates in a flowing bioreactor | yes | (53) | |
| [Non-microbial] Hagfish slime | O | | | yes | (54) | |
| [Non-microbial] Lung sputum from humans with cystic fibrosis | O | | | yes | (55) | |

| Species/Sample type | Microbial biofilm (B), Polymer extract (P), or other (O)? | Lifestyle or environment of organism | Culturing method prior to rheology | G` > G``(within the linear viscoelastic region) | Reference | Comments |
|---|---|---|---|---|---|---|
| [Non-microbial] Salmon skin mucus | O | | Mucus scraped from recently deceased salmon | yes | (56) | |
| *Staphylococcus aureus* | O | skin microbe that is an opportunistic wound pathogen | Liquid culture throughout a growth curve; Strains formed different aggregates in culture | Depends | (57) | (over time, the culture becomes elastic dominant, there are also frequency dependencies for cultures) |
| *Staphylococcus aureus* | O | skin microbe that is an opportunistic wound pathogen | Liquid culture throughout a growth curve; Strains formed different aggregates in culture | Depends | (58) | (frequency and time dependent) |
| [Non-microbial] Mucilage from axenic maize roots | O | | | Depends | (59) | |
| [Non-microbial] Snail adhesive mucus | O | | | yes? | (60) | Strain for frequency sweep not shown, strain sweep provides oscillatory stress and not viscoelastic moduli. In frequency sweep, there is one sample (mucus from snails at 45 degree angle) that has a frequency dependency, the rest are elastic dominant |
| *Klebsiella pneumoniae* | P | lung (?) pathogen | precipitated capsular polysaccharide (from microbes grown in agar medium, then solubilized and precipitated) in 100 mM sodium chloride | depends | (61) | Frequency dependent in linear viscoelastic region, viscous dominant at lower frequencies, crosses over around 10 rad/s |
| [Non-microbial] Mucilage from seed coat | P | | | yes | (62) | |
| *Klebsiella pneumoniae* | P | | precipitated capsular polysaccharide (from microbes grown in agar medium, then solubilized | No | (63) | |

| **Species/Sample type** | **Microbial biofilm (B), Polymer extract (P), or other (O)?** | **Lifestyle or environment of organism** | **Culturing method prior to rheology** | **G` > G``(within the linear viscoelastic region)** | **Reference** | **Comments** |
|---|---|---|---|---|---|---|
| | | | and precipitated) in 100 mM sodium chloride | | | |
| [Non-microbial] Coral mucus | P | | "Milking of corals" method used, then mucus was treated to prevent enzymatic activity. After enzymatic inhibition, a mucin extraction process was carried out | yes (mostly, the data is noisy at low frequencies and there are a few data that show viscous dominance) | (64) | |
| *Bacillus subtilis* γ-polygluconate extract | P | soil bacteria | biofilm grown on agar media | yes (however some conditions of the γ-polygluconate extract were viscous dominant) | (43) | |
| *Azotobacter vinelandii* alginate extract | P | soil bacteria | biofilm grown on agar media | yes | (43) | |
| purified succinoglycans from *Sinorhizobium meliloti* | P | | Grown in glutamate-mannitol-salts liquid media, centrifuged (20,000 g, 5 min), supernatant used for polymer extraction | Depends | (65) | There was both a strain dependency and frequency dependency, amplitude/strain sweep not shown |
| [Non-microbial] Cold-water-extractable mucilage of Plantago ovata seeds | P | | | no | (66) | This is the only biopolymer that has behaved in a G''-dominated manner like Ralstonia biofilms |
| [Non-microbial] Hot water-extractable mucilage of Plantago ovata seeds | P | | | yes | (66) | |
| *Pantoea* sp. YR343 (and it's corresponding ΔUDP mutant) | B | Poplar rhizosphere | Grown in SOBG agar medium (agar, tryptone, yeast extract, NaCl, MgSO4, KCl, & Glycerol) | Yes | (67) | |

**Table S2.** Bacterial strain list.

| Strain | Abbreviation in paper | Isolation source | Genotype | Reference for strain | Lab with Strain | Lab Strain # | Strain obtained from | Acknowledgement |
|---|---|---|---|---|---|---|---|---|
| *Ralstonia pseudosolanacearum* GMI1000 | Rpseu GMI1000 | wilted plant | wild type | (68) | T. Lowe-Power (UC Davis) | UCD1 | C. Allen (UW Madison) | |
| GMI1000 Δ*epsB* | | -- | markerless deletion of RSp1018 created by *sacB* mutagenesis | (11) | T. Lowe-Power (UC Davis) | UCD140 | C. Allen (UW Madison) | |
| GMI1000 Δ*nucAB* | | -- | allelic replacement of *nucA* (RSc0744) with GmR cassette and *nucB* (RSc2452) with KanR cassette | (12) | T. Lowe-Power (UC Davis) | UCD144 | C. Allen (UW Madison) | |
| GMI1000 Δ*lecX* | | -- | markerless deletion of RSp0569 created by *sacB* mutagenesis | (11) | T. Lowe-Power (UC Davis) | UCD583 | C. Allen (UW Madison) | |
| GMI1000 Δ*lecF* | | -- | markerless deletion of RSc2107 created by *sacB* mutagenesis | (11) | T. Lowe-Power (UC Davis) | UCD584 | C. Allen (UW Madison) | |
| GMI1000 Δ*lecFX* | | -- | markerless deletion of RSp0569 and RSc2107 created by *sacB* mutagenesis | (68) | T. Lowe-Power (UC Davis) | UCD585 | C. Allen (UW Madison) | |
| *Ralstonia solanacearum* IBSBF1503 | Rsol IBSBF1503 | wilted plant | Wild type | (69) | T. Lowe-Power (UC Davis) | UCD53 | C. Allen (UW Madison) | |

| Strain | Abbreviation in paper | Isolation source | Genotype | Reference for strain | Lab with Strain | Lab Strain # | Strain obtained from | Acknowledgement |
|---|---|---|---|---|---|---|---|---|
| *Ralstonia syzygii* PSI07 | Rsyz PSI07 | wilted plant | Wild type | (70) | T. Lowe-Power (UC Davis) | UCD5 | C. Allen (UW Madison) | |
| *Xanthomonas campestris* pv. *campestris* 8004 | XCC8004 | XCC8000 was isolated from Brassica oleracea var. botrytis in 1958 | XCC8004 is a Spontaneous rifampicin mutant of Xanthomonas campestris pv. campestris 8000 | (71) | T. Lowe-Power (UC Davis) | UCD611 | P. Ronald (UC Davis) | |
| *Ralstonia insidiosa* 130770013-1 | 130770013-1 | International Space Station | | (72) | Wargo (UVM) | MJ602 | | Acknowledgement for C. Mark Ott, NASA Johnson Space Center |
| *Ralstonia insidiosa* AU13589 | AU13589 | Cystic Fibrosis Sputum | | (73) | Wargo (UVM) | MJ875 | | Acknowledgement for Cystic Fibrosis Stock Center-University of Michigan. |
| *Ralstonia pickettii* AU14824 | AU14824 | Cystic Fibrosis Sputum | | (73) | Wargo (UVM) | MJ876 | | Acknowledgement for Cystic Fibrosis Stock Center-University of Michigan. |
| *Ralstonia pickettii* AU15492 | AU15492 | Deep Wound | | (73) | Wargo (UVM) | MJ877 | | Acknowledgement for Cystic Fibrosis Stock Center-University of Michigan. |
| *Ralstonia insidiosa* AU19393 | AU19393 | Liver Biopsy | | (73) | Wargo (UVM) | MJ885 | | Acknowledgement for Cystic Fibrosis Stock Center-University of Michigan. |
| *Ralstonia pickettii* AU20306 | AU20306 | Cystic Fibrosis Sputum | | (73) | Wargo (UVM) | MJ887 | | Acknowledgement for Cystic Fibrosis Stock Center-University of Michigan. |

| Strain | Abbreviation in paper | Isolation source | Genotype | Reference for strain | Lab with Strain | Lab Strain # | Strain obtained from | Acknowledgement |
|---|---|---|---|---|---|---|---|---|
| *Ralstonia pickettii* AU21292 | AU21292 | Blood | | (73) | Wargo (UVM) | MJ892 | | Acknowledgement for Cystic Fibrosis Stock Center-University of Michigan. |
| *Ralstonia pickettii* AU28504 | AU28504 | Cystic Fibrosis Sputum | | (73) | Wargo (UVM) | MJ899 | | Acknowledgement for Cystic Fibrosis Stock Center-University of Michigan. |
| *Ralstonia pickettii* AU40593 | AU40593 | Endo-tracheal tube aspirate from CF | | (73) | Wargo (UVM) | MJ914 | | Acknowledgement for Cystic Fibrosis Stock Center-University of Michigan. |
| *Ralstonia insidiosa* AU5199 | AU5199 | Blood | | (73) | Wargo (UVM) | MJ916 | | Acknowledgement for Cystic Fibrosis Stock Center-University of Michigan. |
| *Ralstonia pickettii* HI2644 | HI2644 | Soil | | (73) | Wargo (UVM) | MJ920 | | Acknowledgement for Cystic Fibrosis Stock Center-University of Michigan. |
| *Ralstonia pickettii* HI3561 | HI3561 | Environmenta l | | (73) | Wargo (UVM) | MJ922 | | Acknowledgement for Cystic Fibrosis Stock Center-University of Michigan. |
| *Ralstonia pickettii* HI3575 | HI3575 | Environmenta l | | (73) | Wargo (UVM) | MJ923 | | Acknowledgement for Cystic Fibrosis Stock Center-University of Michigan. |
| *Ralstonia pickettii* HI3597 | HI3597 | Environmenta l | | (73) | Wargo (UVM) | MJ924 | | Acknowledgement for Cystic Fibrosis Stock Center-University of Michigan. |

| Strain | Abbreviation in paper | Isolation source | Genotype | Reference for strain | Lab with Strain | Lab Strain # | Strain obtained from | Acknowledgement |
|---|---|---|---|---|---|---|---|---|
| *Ralstonia pickettii* HI3631 | HI3631 | Environmental | | (73) | Wargo (UVM) | MJ925 | | Acknowledgement for Cystic Fibrosis Stock Center-University of Michigan. |
| *Ralstonia insidiosa* HI4151 | HI4151 | Water | | (73) | Wargo (UVM) | MJ926 | | Acknowledgement for Cystic Fibrosis Stock Center-University of Michigan. |
| *Ralstonia thomasii* AB22-23 | AB22-23 | Harvard Forest Long Term Warming Experiment soil, mineral horizon, control treatment | | (74) | DeAngelis (UMass-Amherst) | AB22-23 | | |
| *Ralstonia thomasii* GP71 | GP71 | Harvard Forest Long Term Warming Experiment soil, mineral horizon, control treatment | | (74) | DeAngelis (UMass-Amherst) | GP71 | | |
| *Ralstonia thomasii* GP95 | GP95 | Harvard Forest Long Term Warming Experiment soil, mineral horizon, control treatment | | (74) | DeAngelis (UMass-Amherst) | GP95 | | |
| *Ralstonia thomasii* AB28-3 | AB28-3 | Harvard Forest Long Term Warming Experiment soil, mineral horizon, warming treatment | | (74) | DeAngelis (UMass-Amherst) | AB28-3 | | |

| Strain | Abbreviation in paper | Isolation source | Genotype | Reference for strain | Lab with Strain | Lab Strain # | Strain obtained from | Acknowledgement |
|---|---|---|---|---|---|---|---|---|
| *Ralstonia thomasii* AB36-13C | AB36-13C | Harvard Forest Long Term Warming Experiment soil, mineral horizon, warming treatment | | (74) | DeAngelis (UMass-Amherst) | AB36-13C | | |
| *Ralstonia thomasii* GP101 | GP101 | Harvard Forest Long Term Warming Experiment soil, organic horizon, warming treatment | | (74) | DeAngelis (UMass-Amherst) | GP101 | | |
| *Ralstonia thomasii* vGP103 | GP103 | Harvard Forest Long Term Warming Experiment soil, organic horizon, warming treatment | | (74) | DeAngelis (UMass-Amherst) | GP103 | | |

**Movie S1 (separate file). Colony biofilms of wildtype GMI1000 are fluid enough to drip off their agar plates.** The medium in these petri dishes is CPG broth with 1.5% w/v agar and 0.002% w/v tetrazolium chloride added. Frozen bacterial glycerol stocks are swabbed with a sterile wooden stick and then streaked out onto the agar medium. The plates were incubated at 28 ℃ for 2 days until visible colonies had grown, then plates were incubated at room temperature (~22 °C) for an additional 4 days. These plates were then inverted as seen in the videos to record the ability of the colonies to flow and drip.
